# Engineered odorant receptors illuminate structural principles of odor discrimination

**DOI:** 10.1101/2023.11.16.567230

**Authors:** Claire A. de March, Ning Ma, Christian B. Billesbølle, Jeevan Tewari, Claudia Llinas del Torrent, Wijnand J. C. van der Velden, Ichie Ojiro, Ikumi Takayama, Bryan Faust, Linus Li, Nagarajan Vaidehi, Aashish Manglik, Hiroaki Matsunami

**Author notes:** These authors contributed equally (Claire A. de March, Ning Ma, Christian B. Billesbølle). Correspondence to: Claire A. de March, Nagarajan Vaidehi, Aashish Manglik, or Hiroaki Matsunami.

## Abstract

A central challenge in olfaction is understanding how the olfactory system detects and distinguishes odorants with diverse physicochemical properties and molecular configurations. Vertebrate animals perceive odors via G protein-coupled odorant receptors (ORs). In humans, ∼400 ORs enable the sense of smell. The OR family is composed of two major classes: Class I ORs are tuned to carboxylic acids while Class II ORs, representing the vast majority of the human repertoire, respond to a wide variety of odorants. How ORs recognize chemically diverse odorants remains poorly understood. A fundamental bottleneck is the inability to visualize odorant binding to ORs. Here, we uncover fundamental molecular properties of odorant-OR interactions by employing engineered ORs crafted using a consensus protein design strategy. Because such consensus ORs (consORs) are derived from the 17 major subfamilies of human ORs, they provide a template for modeling individual native ORs with high sequence and structural homology. The biochemical tractability of consORs enabled four cryoEM structures of distinct consORs with unique ligand recognition properties. The structure of a Class I consOR, consOR51, showed high structural similarity to the native human receptor OR51E2 and yielded a homology model of a related member of the human OR51 family with high predictive power. Structures of three Class II consORs revealed distinct modes of odorant-binding and activation mechanisms between Class I and Class II ORs. Thus, the structures of consORs lay the groundwork for understanding molecular recognition of odorants by the OR superfamily.

## Introduction

Vertebrate animals perceive odors primarily through olfactory G protein-coupled receptors (GPCRs) found within sensory neurons of the olfactory epithelium. In humans, olfactory GPCRs account for over half of the class A GPCR family ^1–3^ (**Supplementary Fig. 1a**). Two types of GPCRs are involved in sensing odorants: a large family of odorant receptors (ORs) commonly subdivided further into Class I and Class II types and a separate family of Trace Amine-Associated Receptors (TAARs)^4,5^. Class II ORs are most prevalent, accounting for 84% of all olfactory GPCRs with 335 identified members. They are followed by Class I ORs (56 members) and TAARs (6 members). ORs are further divided into 17 subfamilies (Class II: 1-14; Class I: 51, 52, 56) based on their amino acid sequence similarities^6^.

The olfactory system needs to detect and discriminate odorants with diverse physicochemical properties and molecular structures. This challenging task is accomplished by combinatorial activation of olfactory GPCRs, wherein a single receptor can be activated by multiple odorants and a single odorant can activate multiple receptors^7,8^. Each type of olfactory GPCR is responsible for detecting a particular segment of odor chemical space. While TAARs are specialized to amines and Class I ORs are tuned to carboxylic acids, Class II ORs respond to a much larger array of volatile odorants^9,10^. TAARs and Class I ORs are more abundant in fish, likely because these receptors recognize water soluble odorants. Class II ORs have undergone dramatic expansion in terrestrial vertebrates, likely because they recognize a more diverse set of volatile, poorly water soluble odorants^11,12^. The anatomical distribution of ORs in amphibian species further supports this mapping of chemical diversity to OR classes. In the model amphibian *Xenopus laevis*, Class I ORs are expressed in olfactory epithelium regions dedicated to the detection of water-soluble molecules, while Class II ORs are found in areas dedicated to the detection of volatile odorants^13^.

Several advances have started to provide an atomic perspective on how odorants are recognized by the olfactory system. We recently reported the structure of a human odorant receptor, OR51E2, bound to the odorant propionate^14^. Like most other Class I ORs, OR51E2 responds to carboxylic acids. Additional structural biology studies have reported structures of murine TAARs mTAAR7f^15^ and mTAAR9^16^ bound to linear and cyclic amines. Despite these foundational insights into odorant recognition, how Class II ORs interact with diverse odorants remains elusive for two reasons: 1) Class II ORs share only 18-34% amino acid identity with OR51E2 and 2) Class II ORs recognize a distinct set of odor chemical space compared to Class I ORs and TAARs^9,10^.

To understand how the sequence diversity of ORs enables recognition of diverse odorants, we used a combination of odorant receptor engineering and cryogenic electron microscopy (cryo-EM) to unravel fundamental features of odorant recognition in Class I and Class II ORs. A combination of molecular dynamics simulations and mutagenesis studies revealed key differences in how each of these families recognizes odorants as well as important similarities in how odorants activate their receptors. Furthermore, our engineering strategy to enable structure determination of otherwise technically recalcitrant ORs enables a path to modeling the thousands of ORs encoded across vertebrate genomes.

### Consensus ORs are robustly expressed

A fundamental challenge in the study of vertebrate ORs is low expression levels of native receptors in heterologous cell systems^17^. Our recent structure determination of human OR51E2 relied on identification of an OR that is atypically highly expressed in model cell lines, likely because it is ectopically expressed and strongly conserved during evolution^14^. The vast majority of other vertebrate ORs have remained recalcitrant to overexpression in heterologous cell lines, even with co-expression of dedicated OR chaperones^18–20^. Due to these fundamental challenges in biochemical study of OR function, we applied a previously-established “consensus” strategy for engineering thermostable proteins^21–23^. While initially described for immunoglobulins^24^ and enzymes^25^, we previously demonstrated that consensus OR constructs (consORs) can be designed using individual members of a subfamily of human ORs^26^. Such consORs are expressed in heterologous cells at levels that approach other non-olfactory Class A GPCRs. Importantly, consORs are a robust starting point for modeling individual native ORs as they have high sequence identity to each individual member of an OR subfamily (**Supplementary Table 1**). ConsORs are often activated by similar odorants as their corresponding native ORs.

We initially applied the consensus approach to study the human OR51 subfamily, which belongs to Class I ORs that recognize carboxylic acid odorants. After aligning 23 members of the OR51 subfamily, we designed a consensus construct (consOR51) that retains the most common amino acid at each aligned position (**Fig. 1a**). Phylogenetic analysis of consOR51 compared to the native sequences of OR51 subfamily members shows that the consensus construct lies at the root of the extant sequences, which range from 45% to 74% amino acid identity when compared with consOR51 (**Fig. 1b and Supplementary Table 1)**. The vast majority of individual OR51 subfamily members fail to express at measurable levels in HEK293T cells, with the exception of OR51E2 and, to a lesser extent, OR51E1. By contrast, consOR51 expresses at levels higher than OR51E2 (**Fig. 1c and Supplementary Fig. 2**). In a GloSensor cAMP production assay, consOR51 shows significant elevation of the GloSensor signal at baseline, which suggests that the consensus construct has a high basal activity in the absence of an odorant (**Fig. 2d**).

**Figure 1.**
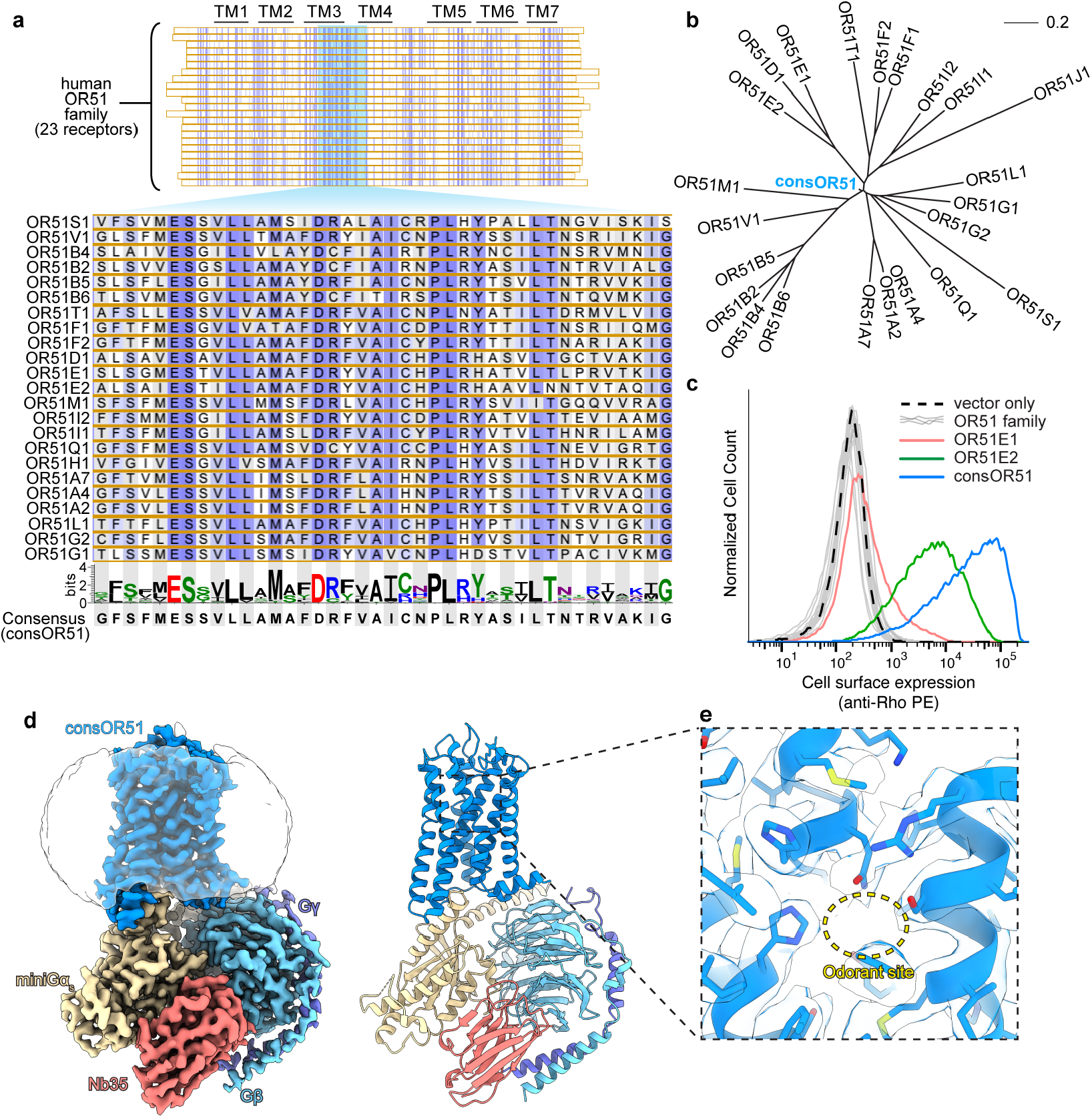
Consensus odorant receptor strategy. **a)** Consensus odorant receptor (consOR) design strategy. All 23 human OR51 subfamily sequences are aligned and the most conserved amino acid is selected at each position to create a consensus sequence. The conserved region in TM3 of the OR51 subfamily is highlighted here. **b)** Phylogenetic tree of the OR51 subfamily including consensus OR51 (consOR51), which occupies the root of the subfamily tree. **c)** Cell surface expression of HEK293 cells transiently transfected with vector control, individual OR51 family members, or consOR51. Most OR51 family members are poorly expressed at the cell surface, with the exception of OR51E2. ConsOR51 shows a dramatic increase in cell surface expression. **d**) Cryo-EM density map of consOR51in complex with G_s_ heterotrimer and stabilizing nanobody Nb35. **e**) Zoom in view of the putative odorant binding site in consOR51 shows a lack of identifiable density for an odorant.

**Figure 2.**
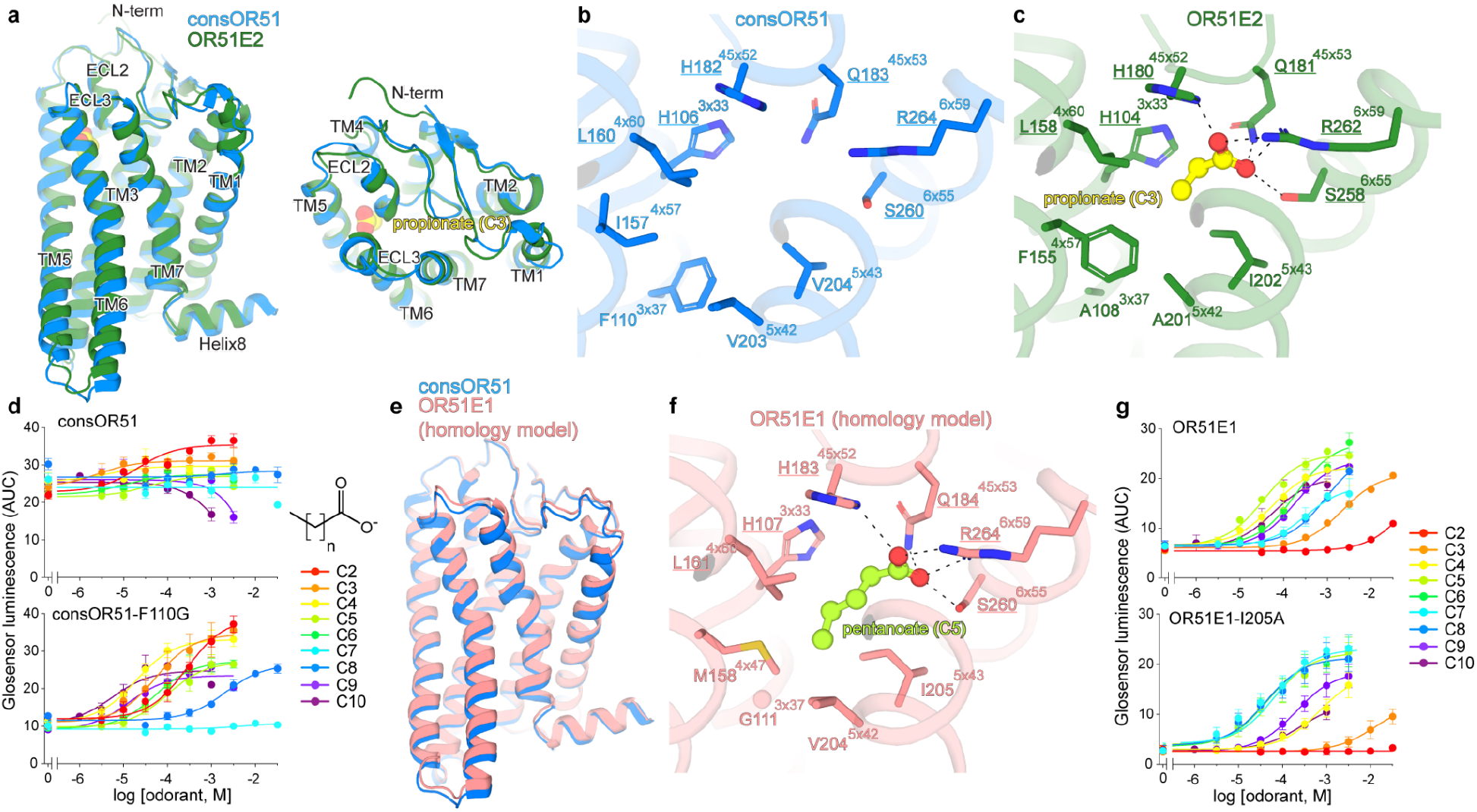
Structure of consOR51 provides insight into native OR51 family members. **a)** Comparison of cryo-EM structure of consOR51 to cryo-EM structure of human OR51E2 indicates high degree of similarity in the 7TM domains and the extracellular loops. Close-up view of odorant binding pocket in consOR51 **(b)** compared to the propionate binding pocket of OR51E2 **(c).** Conserved side chains show similar rotamers. **d**) ConsOR51 is constitutively active in a Glosensor cAMP production assay. Introduction of the F110G mutation in consOR51 leads to suppression of basal activity and response to fatty acids of varying aliphatic chain length. Data points are mean ± standard deviation from n = 3 replicates. **e)** A homology model of human OR51E1 was constructed using consOR51. **f**) Docked structure of pentanoic acid in the OR51E1 homology model. **g)** OR51E1 recognizes long-chain fatty acids, with a preference for pentanoic acid (C5). Selectivity for fatty acid chain length is altered in OR51E1-I205A. Data points are mean ± standard deviation from n = 3 replicates.

Encouraged by the surface expression levels of consOR51, we determined a cryo-EM structure of consOR51. Following the successful strategy used for structure determination of OR51E2, we designed a construct fusing consOR51 with a C-terminal miniGɑ_s_ protein^14,27^. Because consOR51 is constitutively active, we purified the consOR51-miniGɑ_s_ fusion protein in the absence of an odorant agonist. Consistent with increased cell surface expression of consOR51 compared to OR51E2, we observed significantly greater protein purification yields for consOR51-miniGɑ_s_ compared to OR51E2-miniGɑ_s_. We further added Gβ_1_γ_2_ and the stabilizing nanobody Nb35 to produce a complex amenable for single particle cryo-EM studies, which yielded a map of consOR51 bound to the G_s_ heterotrimer with 3.2 Å resolution (**Fig. 1d, Supplementary Fig. 3, and Supplementary Table 2**). Perhaps due to the constitutive activity of consOR51, we did not observe an odorant bound to consOR51 (**Fig. 1e**). Application of the consensus strategy, therefore, enables robust expression of model ORs making them amenable to structure determination.

### Structure of consOR51 enables dissection of OR51 family

We first compared the structures of consOR51 and human OR51E2 to understand how well consensus OR constructs recapitulate the structure of native ORs. The overall structure of consOR51 and OR51E2 are highly similar, with an overall root mean square deviation (RMSD) of 1.3 Å (**Fig. 2a)**. Although the overall architecture of the extracellular loops is highly similar between consOR51 and OR51E2, the intracellular ends of transmembrane helices 5 (TM5) and TM6 deviate slightly between consOR51 and OR51E2. These differences could be due to high basal activity of the consOR51.

A potential utility of consORs is that they may enable accurate modeling of the odorant binding pocket of native ORs. We therefore compared how well consOR51 recapitulates the binding pocket of OR51E2 (**Fig. 2b,c**). Although our structure of consOR51 was obtained without an odorant, comparison of the binding pockets of consOR51 and OR51E2 revealed remarkable similarity in the identity of many amino acids in this region and the conformation of side chains that engage odorants. Perhaps most notable is a conserved arginine residue in Class I ORs (R264^6x59^ in consOR51 and R262^6x59^ in OR51E2, superscripts represent the modified Ballesteros-Weinstein numbering system for GPCRs^28–30)^. We have previously demonstrated that engaging the carboxylic acid of propionate by R262^6x59^ in OR51E2 is critical for receptor activation^14^. In consOR51, we observe that R264^6x59^ is poised to make a similar contact with a carboxylic acid in the odorant binding pocket (**Fig. 2b,c**). More broadly, other residues that engage the propionate carboxylic acid moiety in OR51E2 are similarly poised to interact with a carboxylic acid in consOR51. For OR51E2, we previously demonstrated that hydrophobic interactions between the aliphatic tail of fatty acids and the odorant binding pocket confer fatty acid mediated activity and selectivity. As expected, residues in this region diverge between OR51E2 and consOR51. A notable difference occurs at position 3x37, which is a bulky aromatic in consOR51 (F110^3x37^) compared to a small aliphatic side chain in OR51E2 (A108^3x37^). It has already been shown in a mouse OR that bulky amino acids in this area increase the basal activity of ORs^31^. Mutation of consOR51 at this position to glycine (consOR51-F110G) yielded significantly reduced basal activity and a gain of odorant-dependent response. The increased space at position 4x47 (F155 in OR51E2, I157 in consOR51) accommodates longer chain fatty acids^14^ and, as expected, consOR51-F110G responds best to medium chain length fatty acids (**Fig. 2d**).

We next turned to understand whether the consOR51 structure may enable accurate homology modeling of a different OR51 family member, OR51E1 (**Fig. 2e**). While OR51E2 is selective for the short-chain fatty acids acetate and propionate, OR51E1 responds to longer-chain fatty acids^8,32^. Indeed, in a GloSensor cAMP accumulation assay, we observe that OR51E1 responds to a range of fatty acids, with a preference for pentanoate (pEC50 = -4.64 ± 0.03, **Fig. 2g**). We generated a homology model of OR51E1 using the structure of consOR51 as template, and docked pentanoate into this model (**Fig. 2f**). Similar to the binding pose of propionate in OR51E2, the carboxylic acid of pentanoic acid engages a similar ionic and hydrogen bonding network anchored by R264^6x59^. A distinct set of residues in the divergent part of the cavity enables the longer aliphatic chain of pentanoate to bind in the OR51E1 pocket. To test this model, we mutated residue I205^5x43^ to alanine, predicting that introducing more space in this region would change the fatty acid preference of OR51E1. Indeed OR51E1-I205^5x43^A shows a distinct preference for the longer chain heptanoic and octanoic fatty acids (**Fig. 2g**).

With these studies, we surmise that: 1) consORs likely show high structural similarity to individual native ORs and 2) homology modeling of native ORs from a consOR can enable predictive models of odorant binding.

### Structure of consOR1 as a representative Class II OR

Our structural insights into ORs have thus far been limited to Class I ORs. Attempts to express and purify Class II ORs have been even more challenging than Class I ORs, likely because Class II ORs are generally more poorly folded and induce stronger ER stress responses^33^. Class II ORs recognize a broad range of odorants with significant structural diversity^8,34–36^. Among Class II ORs, the human OR1A1 receptor has previously been characterized as a broadly tuned receptor that recognizes highly diverse odorants, including allyl phenyl acetate, dihydrojasmone, menthols, and carvones^37^. We, therefore, sought to understand how the binding pocket of OR1A1 leads to its specific odorant recognition profile.

We started by using the consensus approach to generate consOR1, a construct that shares 63% sequence identity with native OR1A1 (**Fig. 3a)**. In contrast to consOR51, consOR1 is not constitutively active and responds robustly to the odorant L-menthol (**Fig. 3b,c, and Supplementary Fig. 5**). Like OR1A1, consOR1 responds to a diverse set of odorants, highlighting the unique ability of consensus ORs to recapitulate features of native ORs. Using a similar strategy as for consOR51, we determined a cryo-EM structure of consOR1 bound to L-menthol with a nominal resolution of 3.3 Å (**Fig. 3d, Supplementary Fig. 4, and Supplementary Table 2**).

**Figure 3.**
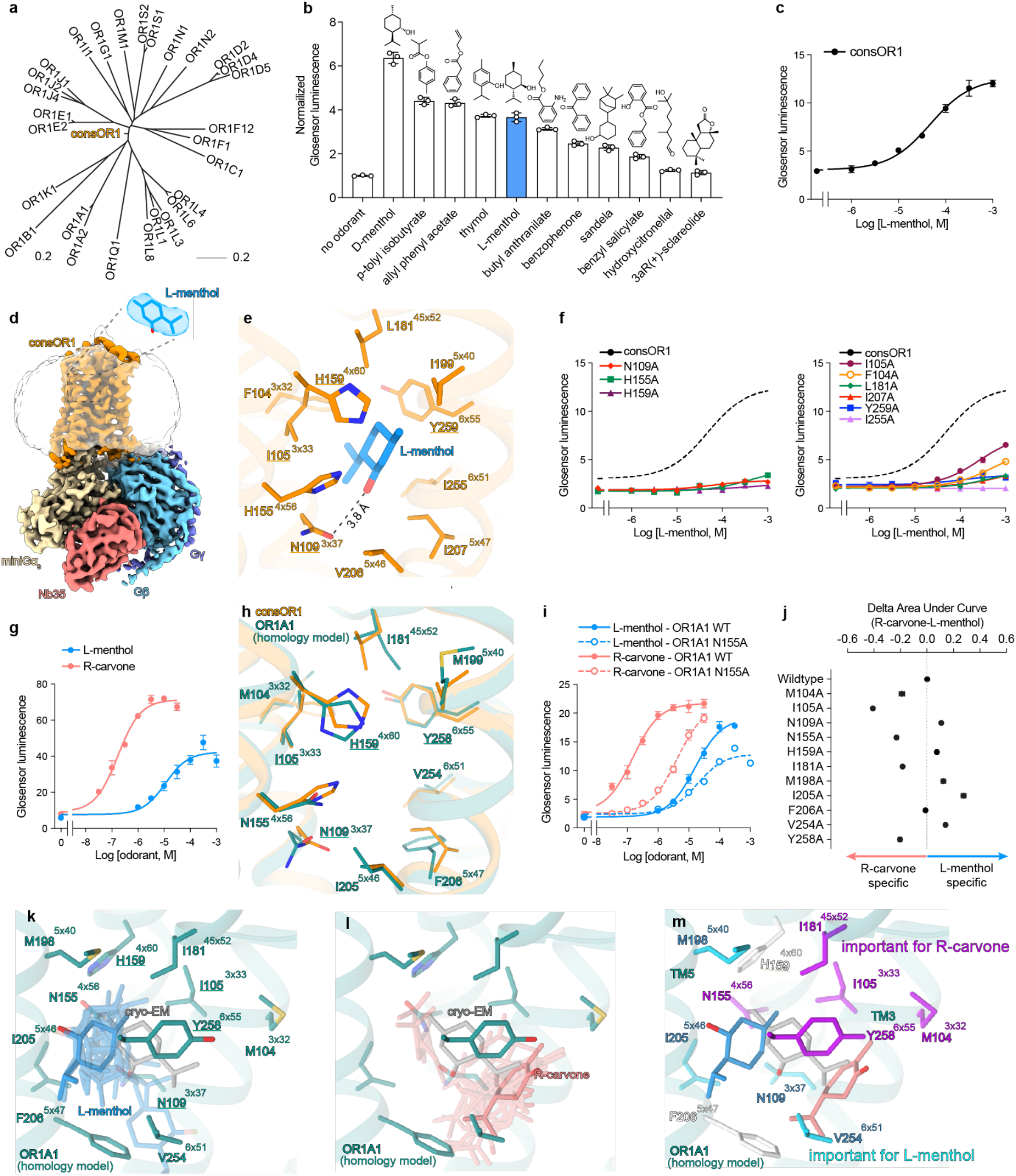
The structure of consOR1 provides insight into human OR1A1. **a)** Phylogenetic tree of the human OR1 subfamily including consOR1. **b)** ConsOR1 is activated by diverse odorants as measured by a Glosensor cAMP production assay. Area under the dose response curve was calculated and normalized to the no odorant negative control (n = 3). **c)** Dose response for L-menthol activation of consOR1. **d)** Cryo-EM map of the consOR1-G_s_ complex. Insert shows cryo-EM density for L-menthol. **e)** View of the consOR1 odorant binding pocket within 5 Å with a single hydrogen bond shown as dashed lines. **f)** Mutagenesis studies of consOR1 in a cAMP accumulation assay. **g)** OR1A1 is activated by terpenoids L-menthol and R-carvone. **h)** Homology model of OR1A1 based on consOR1 **i)** The OR1A1-N155A mutation has a larger effect on R-carvone activity as compared to L-menthol. **j)** OR1A1 mutants differentially affect R-carvone or L-menthol activity. Area under the dose-response curve was calculated for each OR1A1 mutant activated by either odorant (n = 3). For each odorant, AUC values were normalized to wildtype OR1A1. Subtraction of normalized AUCs revealed a differential effect of mutations. Docking of L-menthol (**k**) and R-carvone (**l**) docked to the homology model of OR1A1. Top scoring docking results are shown for both odorants as transparent sticks. The best scoring pose is shown as solid sticks. **m)** Mapping the effect of mutations in (j) onto the homology modeled structure of OR1A1 shows that mutations that affect L-menthol and R-carvone cluster in distinct regions of the OR1A1 binding pocket. Every residues with a delta of more than 0.10 are reported colored by the odorant the most affected. For all cell assay, data points are mean ± standard deviation from n = 3 replicates.

The binding pocket of consOR1 is largely hydrophobic with a few amino acids that provide either hydrogen bond donors or acceptors. Cryo-EM density for L-menthol supported a binding pose with the hydroxyl group of the odorant engaging N109^3x37^ in the binding pocket (**Fig. 3e**). L-menthol makes van der Waals contacts with many residues in the consOR1 binding pocket. Indeed, alanine mutagenesis experiments show that the majority of residues within 5 Å from the ligand in the binding pocket are important for L-menthol activity at consOR1 (**Fig. 3f**). We conclude that most residues in the consOR1 binding pocket contribute to L-menthol binding and efficacy. It is likely that many other Class II ORs show similar modes of odorant recognition – a combination of many distributed hydrophobic contacts combined with a limited set of hydrogen-bonding interactions.

### ConsOR1 enabled dissection of a native Class II OR

We next turned to understand odorant recognition by the native receptor OR1A1. Like consOR1, OR1A1 responds to L-menthol with micromolar potency (pEC_50_ = -4.79 ± 0.05, **Supplementary Fig. 5**). We additionally identified several other odorants with activity at OR1A1, and focused on molecular recognition of another terpenoid odorant, R-carvone (**Fig. 3g**). Compared to L-menthol, R-carvone is more potent at OR1A1 (pEC_50_ = -6.82 ± 0.05, **Supplementary Fig. 5**). Both L-menthol and R-carvone are primarily hydrophobic ligands but harbor a single hydrogen bond donor or acceptor. To understand how OR1A1 recognizes these distinct terpenoids, we generated a homology model of OR1A1 based on the structure of consOR1 bound to L-menthol (**Fig. 3h**). This model was used for docking studies of L-menthol and R-carvone. In both cases, we found that docking did not identify a single pose of the odorant within the OR1A1 binding pocket (**Fig. 3k,l**). Instead, both L-menthol and R-carvone dock to OR1A1 in multiple orientations with a distributed set of van der Waals contacts. Despite the shared terpenoid scaffold of both odorants, docking revealed that L-menthol and R-carvone engage distinct subpockets in OR1A1 that are different from the position of L-menthol bound to consOR1 in the cryo-EM structure. In OR1A1, L-menthol engages residues in TM5 more extensively, while R-carvone engages the other side of the pocket composed primarily of residues in TM3 (**Fig. 3m**).

To test these docking predictions, we assessed the activity of L-menthol and R-carvone against alanine mutants of each binding pocket residue (**Fig. 3j, Supplementary Fig. 5**). We anticipated that these mutations may differentially affect the activity of L-menthol and R-carvone due to their distinct engagement of the OR1A1 pocket. Two mutations, F206^5x47^A and H159^4x60^A, are deleterious for both L-menthol and R-carvone activity (**Supplementary Fig. 5**). Other mutations more selectively affect either L-menthol or R-carvone activity. For example, N155^4x56^A leads to a ∼27-fold worse EC_50_ for R-carvone. By contrast, the same mutation has a negligible effect on potency and ∼30% reduced E_max_ for L-menthol (**Fig. 3i**). To more easily capture the combined effects of efficacy and potency, we calculated the integrated area under the dose-response-curve for each mutant (see Methods). Comparison of this metric for L-menthol and R-carvone revealed that OR1A1 mutations have differential effects on the activity of both odorants (**Fig. 3j**). Concordant with our docking analysis, OR1A1 binding pocket mutations in TM3 more strongly affect R-carvone activity, while mutations in TM5 more strongly affect L-menthol activity (**Fig. 3m**).

These docking and mutagenesis studies highlight the complex mode of odorant recognition for a broadly tuned Class II OR, which likely involves many different odorant binding poses. Different odorants likely engage a single odorant receptor binding pocket in distinct ways, further adding complexity to molecular recognition in the odorant receptor system.

### Structural flexibility in consOR1 ligand recognition

The flexibility of R-carvone docking to OR1A1 and site-directed mutagenesis data suggest that odorants can bind Class II ORs without a single, well defined binding pose. Our previous studies of OR51E2 showed that propionate is not flexible in its binding site and persistently adopts a single pose that is constrained by an ionic interaction. Compared to highly water soluble Class I odorants like propionate, Class II OR ligands are more hydrophobic, often with only a single hydrogen bond donor or acceptor (**Supplementary Fig. 1a**). We therefore sought to understand the structural dynamics of odorant binding to Class II ORs.

We turned to all-atom molecular dynamics simulations (MD) to examine the flexibility of L-menthol in the consOR1 binding pocket. To understand how the G protein and ligand influence consOR1 flexibility, we performed simulations under the following conditions: 1) consOR1 bound to L-menthol and miniGα_s_, 2) consOR1 bound to L-menthol without miniGα_s_, and 3) consOR1 alone (**Fig. 4a**). Each simulation was performed in 5 replicates and each replicate was evolved over 1 µs (**Supplementary Fig. 6**). As expected based on simulations of other GPCRs^38–40^, removal of miniG_s_ and L-menthol leads to increased structural flexibility of consOR1 (**Fig. 4a**). Notably, L-menthol is highly dynamic within the ligand binding pocket of consOR1 (**Fig. 4b-d**). In the absence of G protein, L-menthol explores a broad range of the odorant binding site with a ligand RMSD of 6.1 Å when compared to the cryo-EM structure of L-menthol bound to consOR1. In simulations of consOR1 bound to miniGα_s_, the flexibility of L-menthol is reduced, with a ligand RMSD of 4.2 Å. The flexibility of L-menthol stands in stark contrast to the relative stability of propionate bound to OR51E2 (**Fig. 4e-g**). Our previous simulations of propionate bound to OR51E2 revealed an overall ligand RMSD of 2.1 Å and 2.4 Å for simulations performed with and without the miniGα_s_, respectively. The increased flexibility of L-menthol in simulations of consOR1 without miniGα_s_ is correlated with an increase in the volume of the consOR1 binding pocket. With miniGα_s_, L-menthol explores a consOR1 pocket that encloses 250 Å^3^. In the absence of miniGα_s_, the pocket expands to 450 Å^3^ (**Supplementary Fig. 7**). The increased volume of the consOR1 ligand binding pocket arises from an outward movement of extracellular loop 3 and the extracellular sides of TM6 and TM7.

**Figure 4.**
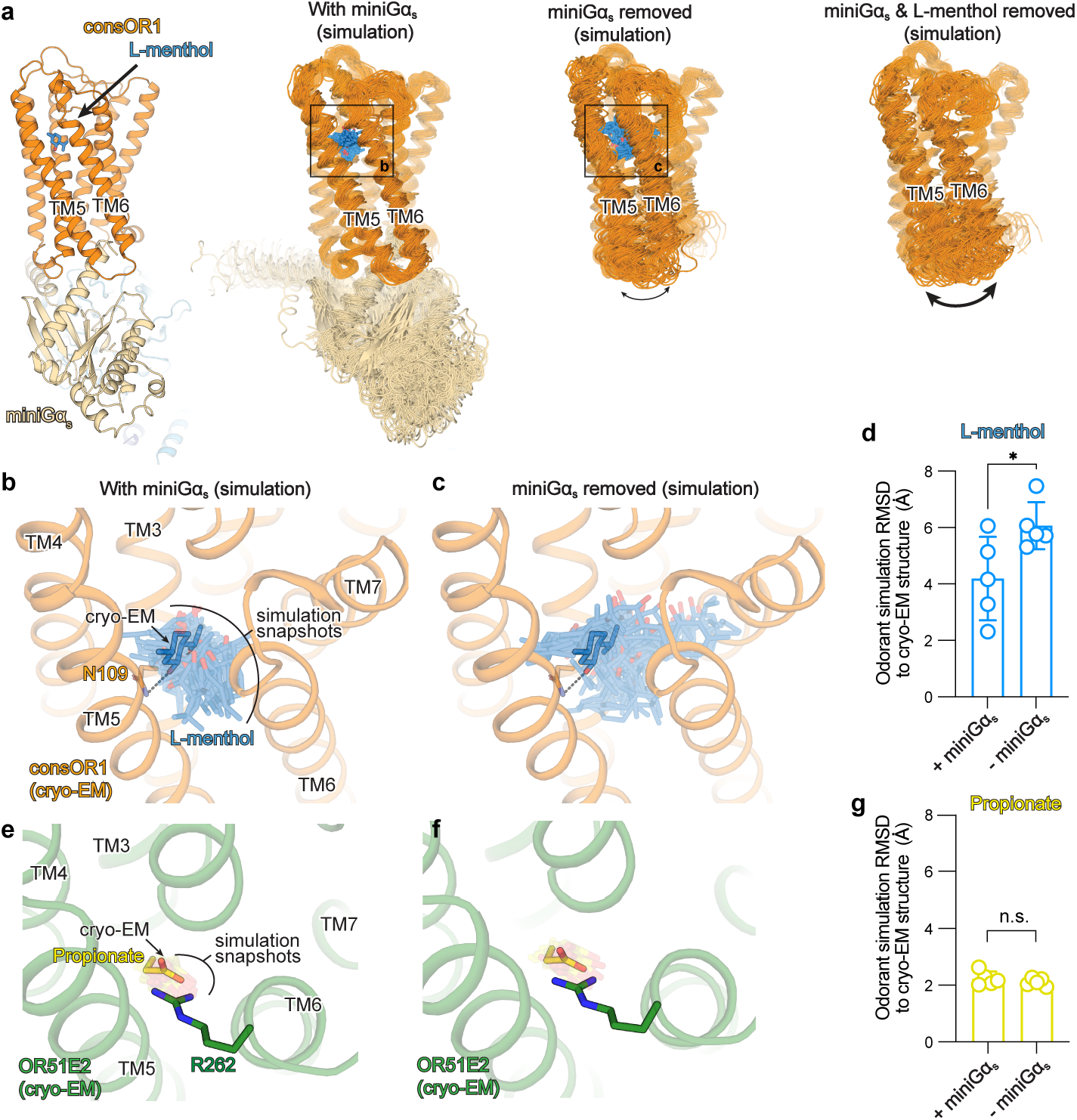
Structural flexibility in odorant binding. **a)** Molecular dynamics simulations were performed for consOR1: with miniGαs and L-menthol, without miniGα_s_ but with L-menthol, and without both miniGα_s_ and L-menthol. Snapshots from 100 ns intervals are shown from a representative simulation. In consOR1, TM5 and TM6 are more flexible in the absence of miniGαs, and even more dynamic in the absence of miniGαs and L-menthol. L-menthol is dynamic in the binding pocket of consOR1 in simulations with miniGα_s_ (**b)**, and shows even greater flexibility in simulations without miniGα_s_ (c). **d)** The root mean squared deviation (RMSD) of L-menthol compared to the cryo-EM pose for each simulation replicate is shown. * indicates p<0.05. **e,f)** In simulations of OR51E2, propionate is constrained within the ligand binding pocket and makes a persistent interaction with R262. **g)** The root mean squared deviation (RMSD) of propionate compared to the cryo-EM pose for each simulation replicate is shown. n.s. Indicates not significant.

Taken together, these simulation and mutagenesis studies suggest that odorants bind to Class II ORs with significantly greater flexibility compared to Class I OR. Furthermore, our simulations also show that binding of the G protein decreases odorant flexibility in a Class II OR binding pocket.

### A shared Class II OR activation motif

We next sought to expand the consOR strategy to other Class II ORs, with the aim of understanding both shared and distinct features between Class I and Class II ORs. We therefore applied the consensus strategy to other human Class II OR subfamilies: the OR2 family (68 members) and the OR4 family (51 members) as shown in **Supplementary Fig. 1**. ConsOR2 and consOR4 respond to the odorants S-carvone and 2-methyl thiazoline (2MT), respectively (**Fig. 5a,b**). We determined cryo-EM structures of consOR2 and consOR4 at 3.2 Å and 3.5 Å, respectively (**Supplementary Figs. 8-9, Supplementary Table 2**). For both receptors, we could identify clear cryo-EM density for the odorant molecules (**Supplementary Fig. 10**). Similar to consOR1, our simulations of consOR2 and consOR4 revealed significant flexibility in the binding pose of odorants at these receptors (**Supplementary Fig. 10**)

**Figure 5.**
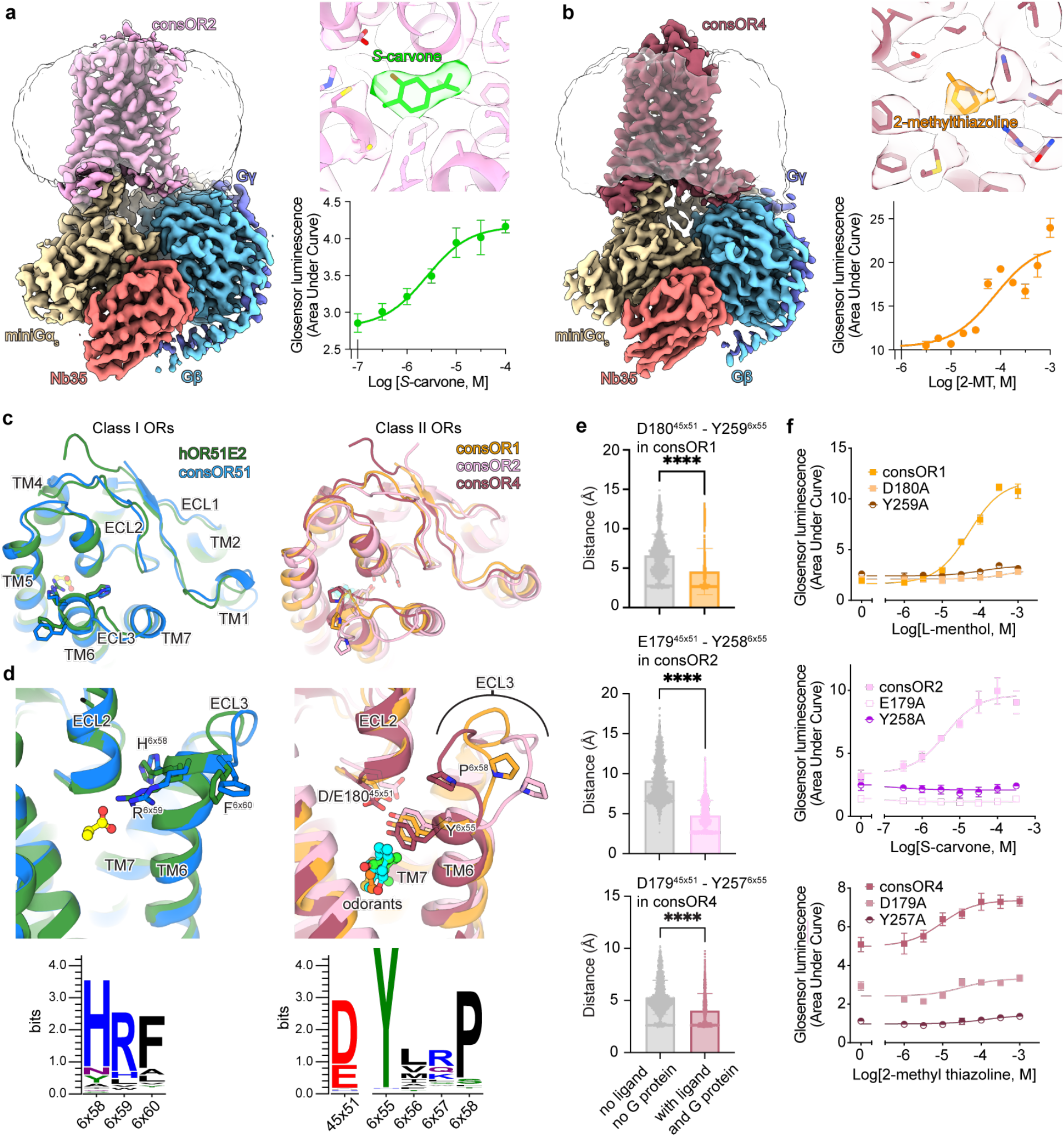
Structures of consOR2 and consOR4 and insights into common features of OR function. **a)** CryoEM map of consOR2-G_s_ complex bound to activating odorant *S*-carvone. **b)** CryoEM map of consOR4-G_s_ complex bound to activating odorant 2-methylthiazoline (2-MT). **c)** Comparison of Class I and Class II OR structures in the extracellular region. Consensus OR structures of Class II ORs show variability in ECL3 conformation. **d)** Close-up view of ligand binding sites in Class I and Class II ORs. Class I ORs recognize carboxylic acids via the R6x59 residue in the extracellular portion of TM6. Class II ORs bind ligands via a highly conserved Y6x55 residue that further engages a conserved acidic residue in ECL2 (D/E^45x^^51^). **e)** The interaction between D/E^45x^^51^ and Y^6x55^ is maintained in simulations of consOR1, consOR2 and consOR4 bound to their agonist and miniGα_s_. Removal of miniGα_s_ and agonist leads to an increase in distance between these two positions (unpaired t-test, p < 0.0001). **f)** Mutation of D^45x51^ and Y^6x55^ in consOR1, consOR2 and consOR4 reduces OR response to odorant in a cAMP production assay. For all cell assay, data points are mean ± standard deviation from n = 3 replicates.

With this set of OR structures, we aimed to identify structural features that are unique to Class I and Class II ORs. As expected, the intracellular regions of Class I and Class II ORs are conserved, both in sequence and structure because these regions are critical for G protein coupling in response to odorant binding (**Supplementary Fig. 11**). The overall fold of the extracellular region is similar between Class I and Class II ORs (**Fig. 5c**).

Despite these similarities, our structural analysis highlighted a common motif in the extracellular region of Class II ORs that is distinct from Class I ORs and is likely important for receptor activation. For OR51E2, we previously demonstrated that a highly conserved arginine residue in Class I ORs (R^6x59^) at the extracellular tip of TM6 engages the carboxylic acid group of fatty acids. This interaction restrains an otherwise dynamic extracellular loop 3 (ECL3), which is associated with receptor activation (**Fig. 5d**). In structures of Class II consORs, position 6x59 is not conserved. Instead, we identified a highly conserved tyrosine residue in Class II ORs (Y^6x55^) that makes a hydrogen-bonding contact with another Class II-specific conserved acidic residue in ECL2 (D/E^45x51^). This conserved contact is in proximity to the odorant binding site, which suggests that it might have an important role in connecting odorant binding to receptor activation. To explore this possibility, we used molecular dynamics simulations to examine this conserved Class II OR contact. With the odorant and G protein bound, this contact is maintained in most simulations across all three Class II consORs. In simulations without the odorant and G protein, the interaction between Y^6x55^ and D/E^45x51^ is less stable, with significantly greater distances over the simulation timeframes (**Fig. 5e)**. Indeed, for all three consORs, disruption of this interaction by alanine mutation markedly reduces odorant-induced activity (**Fig. 5f**). We therefore conclude that the conserved interaction between Y^6x55^ and D/E^45x51^ is an important mechanism for odorant-induced activation of Class II ORs.

## Discussion

Our studies of several OR structures and their dynamic movements yield an emerging general model for odorant recognition. Class I ORs recognize carboxylic acids via a conserved arginine residue in TM6 (R^6x59^). The structure of constitutively active consOR51 captured without an odorant underscores that this residue occupies a conserved position in the binding pocket of activated Class I ORs. Predictive homology modeling of OR51E1 based on consOR51 further supports the following model for class I OR odorant recognition: conserved binding pocket residues that engage the carboxylic acid combined with more divergent binding pocket residues that tune the response profile for fatty acids of varying aliphatic length. Together, these interactions stably position an odorant in the binding site. While odorants bind to the Class II ORs in a similar location as Class I ORs, our studies suggest several distinct mechanisms of odorant recognition between Class I and Class II ORs. First, Class II OR do not harbor a conserved interaction partner analogous to R^6x59^ in class I ORs. Second, odorants make a diffuse set of van der Waals contacts in the Class II OR binding pocket, often with a single hydrogen bonding interaction. For broadly tuned Class II ORs, different odorants are likely to occupy different subpockets of the odorant binding site leading to distinct sets of interactions important for their activity. Third, our studies with consOR1 suggest that odorants bind with significant flexibility in Class II ORs compared to Class I ORs. This likely arises from the more limited set of strong ionic or hydrogen bond contacts in most volatile odorants that activate Class II ORs as compared to the charged water-soluble odorants that activate Class I ORs. An additional factor is likely the increased flexibility of the OR binding pocket in Class II ORs. Recognition of odorants by Class II ORs is therefore also distinct from TAARs, which recognize aminergic odorants via conserved ionic interactions^41^.

More broadly, the vast majority of small molecule binding class A GPCRs use specific hydrogen bonding or ionic interactions to drive specificity in ligand binding. Class II ORs, by contrast, recognize odorants primarily by van der Waals contacts, with limited hydrogen bonding interactions. Our model for odorant recognition in vertebrate ORs recapitulates recent structural biology studies that have identified a flexible binding mode for odorants at a broadly tuned ionotropic insect olfactory receptor, MhOR5^42,43^. In both cases, odorant binding is not confined to a single pose. Despite this flexibility, distinct interactions made between the odorant and OR binding pocket can still result in different odorant activity, as outlined by our studies with R-carvone and L-menthol acting at OR1A1. These distinct sets of interactions drive odorant discrimination. While our studies start to explain some features of molecular recognition in Class II ORs, a more complete understanding how the large diversity of odorants is recognized by this set of ORs will require significant further structural interrogation of both broadly and narrowly tuned receptors.

Our structural analysis also sheds light on a unifying mechanism of Class I and Class II OR activation by chemically diverse odorants. While the specific motifs that engage odorants are distinct between Class I and Class II ORs, a highly conserved interaction between the extracellular end of TM6 and the odorant or odorant binding pocket stabilizes an inward movement of TM6. For Class I ORs, this interaction is driven by odorant engaging R^6x59^ (**Fig. 6a**). For Class II ORs, odorants stabilize an interaction between Y^6x55^ and D/E^45x51^ (**Fig. 6b**). Odorant binding in both Class I and Class II ORs causes an inward movement of the extracellular region in TM6. This movement is accompanied by outward movement of the intracellular side of TM6, which creates a cavity for engaging G protein. Odorants can be structurally flexible while bound to Class II ORs. Full activation of the OR with odorant and G protein restrains some of this flexibility. While a more accurate model will require an experimental structure of an inactive OR, our proposed model provides a shared activation mechanism for the broader OR family.

**Figure 6.**
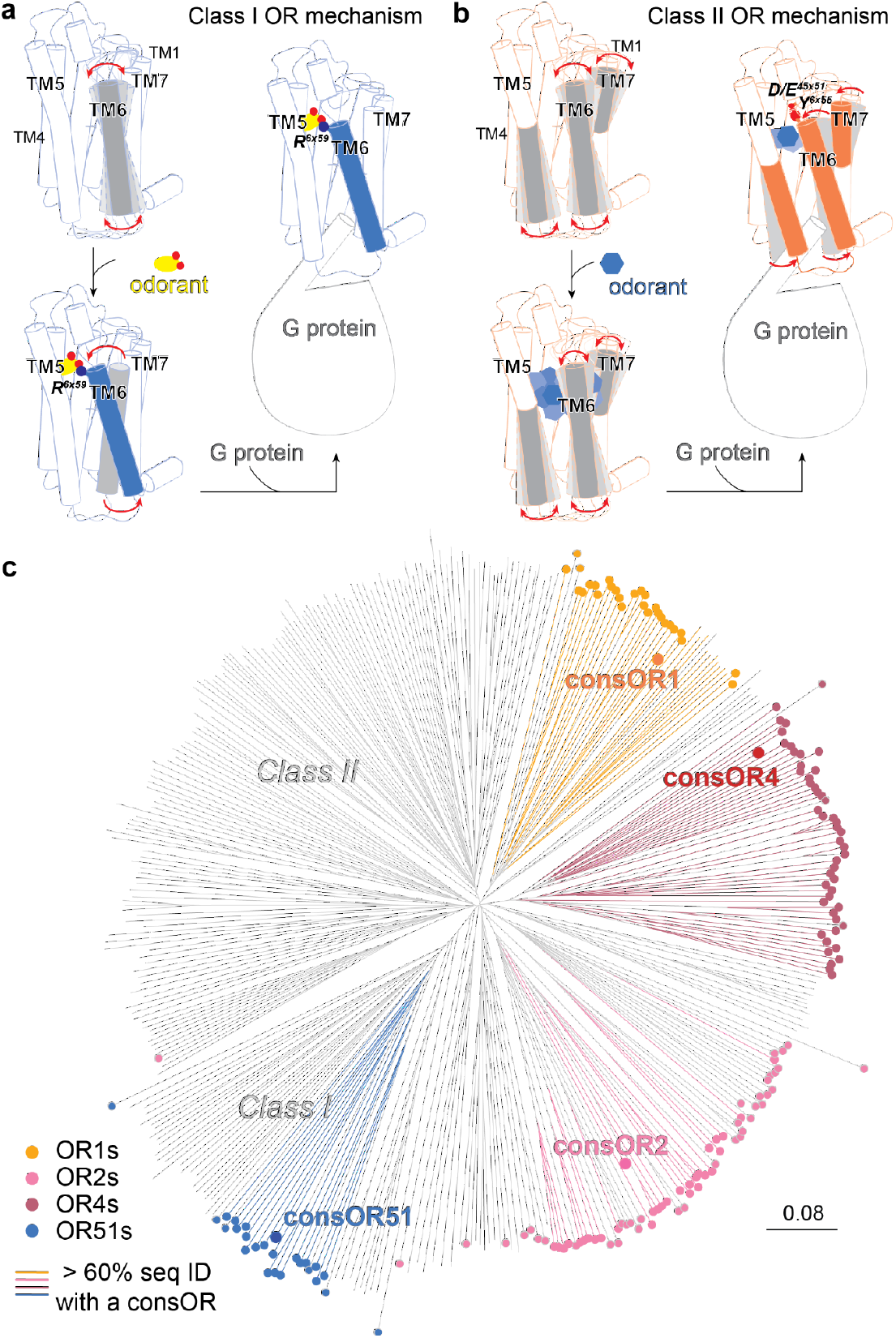
Accessing Class I and Class II OR mechanisms and structures. **a)** Model of Class I OR activation mechanism. **b)** Model of Class II OR activation mechanism. **c)** OR phylogenetic tree including the structurally elucidated consORs showing that these structures allow the homology modeling of 34% of the human native ORs. OR belonging to a consOR subfamily are highlighted by a rounded tip and ORs showing at least 60% of sequence identity with a consOR are shown in colored lines. The scale represents the amount of amino acid change for a set distance.

A key advance of this study is the broad utility of a consensus engineering approach to understand OR function^26^. The vast majority of ORs, in both vertebrate and invertebrate species, remain intractable for biochemical and structural studies. With the consensus approach, we obtained four cryo-EM structures of consORs with high sequence identity to a subset of native human ORs. Our comparison of consOR51 to native human OR51E2 highlights that consOR structures not only share virtually identical backbone structures to native OR family members, but also key residue positions in the structures relevant for odorant recognition. While AlphaFold has transformed protein structure prediction^44^, key OR regions critical for odorant recognition (e.g. ECL3) remain poorly predicted, likely because they diverge so widely across the OR family^14^. We therefore propose that consOR structures will enable higher quality models of many ORs, used either as templates for AlphaFold or in more classic homology modeling approaches. If we use a threshold of 60% sequence identity as a metric for high quality templates for such modeling^45^, the four consOR structures described here would enable high quality models for 34% of all human ORs (**Fig. 6c and Supplementary File 1**). Additional consOR structures derived from the other major OR families will expand this number further. The ability to capture structures of odorants bound to consORs will likely continue to provide fundamental insights into how vertebrate ORs cope with the immense chemical diversity of odorous molecules.

What can the success of consORs reveal about the evolution of the OR family? We previously proposed that the stability of consORs suggests that ancestral OR sequences were likely more stable than the majority of extant OR sequences^26^. Diversification of OR sequences is enabled by evolutionary capacitance provided by OR-specific chaperones^46–48^. The high structural similarity between a consOR and a native OR suggests that evolution drives OR diversity within a family primarily by altering contacts with odorants, as opposed to dramatic variation in the overall fold of the OR. This diversity re-tunes odorant specificity. Furthermore, the fact that the consensus strategy yields stable ORs for multiple OR subfamilies suggests that the common ancestor of each major human OR subfamily was likely a more stable receptor - evolution drives diversification for odorant recognition function at the cost of stability. We anticipate that future studies in visualizing OR structures and odorant recognition will yield deeper insight into the importance of such tradeoffs.

## METHODS

### Expression and purification of consOR-miniG_s_ protein complexes

Expression and purification of consensus OR constructs was done similarly to OR51E2-miniG_s_^14^. Briefly, consensus OR sequences^26^ were cloned into pCDNA-Zeo-TetO with an N-terminal influence hemagglutinin signal sequence and a FLAG (DYKDDDDK) epitope. The construct included the miniG_s399_ protein^27^ fused to the C terminus with a human rhinovirus 3C protease cleavage site. The resulting constructs were transfected into inducible Expi293F-TetR cells using the ExpiFectamine 293 Transfection Kit per the manufacturer’s instructions. After 16 hours, protein expression was induced with 1 µg/mL doxycycline hyclate and the culture was incubated for 36 hours in a shaking incubator maintained at 37 °C and a 5% CO_2_ atmosphere prior to cell harvest by centrifugation.The resulting pellet was stored at -80 °C until purification.

Odorant receptor purification was performed as described previously^14^. Cells pellets were thawed with hypotonic lysis in 20 mM HEPES, pH 7.5, 1 mM EDTA, 100 μM tris(2-carboxyethyl)phosphine (TCEP; Fischer Scientific) and one EDTA-free Protease Inhibitor Tablet (Pierce; ThermoScientific) for 10 min at 4 °C. The lysis buffer was supplemented with odorants to stabilize the consOR constructs: 3 mM L-menthol, 1 mM S-carvone and 30 mM 2-methylthiazoline were used for consOR1-miniG_s399_, consOR2-miniG_s399_ and consOR4-miniG_s399_ purification, respectively. Lysed cells were harvested by centrifugation at 16,000xg for 15 min and immediately dounce-homogenized in ice-cold solubilization buffer comprising 50 mM HEPES, pH 7.5, 300 mM NaCl, 1% (w/v) lauryl maltose neopentyl glycol (L-MNG; Anatrace), 0.1% (w/v) cholesteryl hemisuccinate (CHS, Steraloids), 5 mM adenosine 5′-triphosphate (ATP; Fischer Scientific), 2 mM MgCl2, and 100 μM TCEP. For consOR2-miniG_s399_ and consOR4-miniG_s399_, the solubilization buffer was supplemented with 1 mM S-carvone and 30 mM 2-methylthiazoline, respectively. Due to the low solubility of L-menthol in aqueous buffers, we generated L-menthol doped L-MNG micelles for consOR1-miniG_s399_ purification with a ratio of 0.4 mol% L-menthol in 1% w/v L-MNG. This solution was used in place of 1% L-MNG during purification steps. Solubilized cells were stirred for 1 hour at 4 °C, and the detergent-solubilized fraction was clarified by centrifugation at 16,000xg for 30 min. The detergent-solubilized sample was supplemented with 5 mM CaCl_2_ and incubated in batch with homemade M1-FLAG-antibody conjugated CNBr-Sepharose under slow rotation for 1.5 h at 4 °C. The Sepharose resin was transferred to a glass column and washed with 15 column volumes of ice-cold buffer comprising 50 mM HEPES, pH 7.5, 300 mM NaCl, 0.05% (w/v) L-MNG or 0.05% (w/v) L-MNG with 0.02mol% L-menthol, 0.001% (w/v) CHS, 2.5 mM ATP, 4 mM CaCl_2_, 2 mM MgCl_2_, 100 μM TCEP and the corresponding odorant. This was followed by 10 column volumes of ice-cold 50 mM HEPES, pH 7.5, 150 mM NaCl, 0.0075% (w/v) L-MNG or 0.0075% (w/v) L-MNG with 0.003mol% L-menthol, 0.0025% glyco-diosgenin (GDN; Anatrace), 0.001% (w/v) CHS, 4 mM CaCl2, 100 μM TCEP and corresponding odorant. Receptor-containing fractions were eluted with ice-cold 50 mM HEPES, pH 7.5, 150 mM NaCl, 0.0075% (w/v) L-MNG or 0.0075% (w/v) L-MNG with 0.003mol% L-menthol, 0.0025% (w/v) GDN, 0.001% (w/v) CHS, 5 mM EDTA, 100 μM TCEP, corresponding odorant and 0.2 mg ml−1 FLAG peptide. Fractions containing the consOR–miniG_s399_ fusion protein were concentrated in a 50-kDa MWCO spin filter (Amicon) and further purified over a Superdex 200 Increase 10/300 GL (Cytiva) size-exclusion chromatography (SEC) column, which was equilibrated with 20 mM HEPES, pH 7.5, 150 mM NaCl, 0.0075% (w/v) L-MNG or 0.0075% (w/v) L-MNG with 0.003mol% L-menthol, 0.0025% (w/v) GDN, 0.001% (w/v) CHS, 100 μM TCEP and corresponding odorant. Fractions containing monodisperse consOR–miniG_s399_ were combined and concentrated in a 50-kDa MWCO spin filter.

Other components of the G protein complex, including Gβ_1_γ_2_ and Nb35 were purified as described previously^14,49^. To prepare active-state complexes for cryo-EM, a 3-fold molar excess or 6-fold molar excess of Gβ_1_γ_2_ and Nb35 was added to concentrated consOR4-miniG_s399_ or consOR1-miniG_s399_ and consOR2-miniG_s399_ samples respectively. The resulting preparation was incubated overnight on ice. Complexed samples were purified using a Superdex 200 Increase 10/300 GL SEC column in a buffer comprised of 20 mM HEPES, pH7.5, 150 mM NaCl, 0.0075% (w/v) L-MNG or 0.0075% (w/v) L-MNG with 0.003mol% L-menthol, 0.0025% GDN, and 0.001% w/v CHS, 100 μM TCEP and corresponding odorant. Fractions containing the consOR-G protein complex were collected and concentrated on a 100 kDa MWCO spin filter immediately prior to cryo-EM grid preparation.

### Cryo-EM vitrification, data collection, and processing

The purified OR–G_s_ complex was applied to glow-discharged 300 mesh R1.2/1.3 UltrAuFoil Holey gold support films (Quantifoil). Support films were plunge-frozen in liquid ethane using a Vitrobot Mark IV (Thermo Fisher) with a 10-s hold period, blot force of 0, and blotting time varying between 1.5 and 3 s while maintaining 100% humidity and 4 °C. Vitrified grids were clipped with Autogrid sample carrier assemblies (Thermo Fisher) immediately before imaging. Movies were recorded using a Titan Krios Gi3 (Thermo Fisher) with a BioQuantum Energy Filter (Gatan) and a K3 Direct Electron Detector (Gatan). Data were collected using SerialEM 3.8^50^ running a 3 × 3 image shift pattern at 0° stage tilt. A nominal magnification of ×105,000 with a 100-µm objective was used in super-resolution mode with a physical pixel size as indicated in Supplementary Table 2. Movies were recorded using dose-fractionated illumination with a total exposure of 50 e^−^ Å^−2^ over 60 frames yielding 0.833 e^−^ Å^−2^ per frame. Movies were motion-corrected and Fourier-cropped to physical pixel size using UCSF MotionCor2^51^. Dose-weighted micrographs were imported into cryoSPARC v4.0.3 (Structura Biotechnology^52^), and contrast transfer functions (CTFs) were calculated using the patch CTF estimation tool. Where indicated (see Extended Data Fig. 3, 4, 8, or 9), a threshold of CTF fit resolution was used to exclude low-quality micrographs. Particles were template picked using a 20 Å low-pass-filtered model that was generated ab initio from data collected on the consOR51 sample (Extended Data Fig. 3). Particles were extracted with a box size of 288 pixels, binned, and sorted by 3D classification with alignment using the heterogeneous refinement tool. Template volumes for each of the four classes were low-pass filtered to 20 Å. The resulting particles were re-extracted with a box size of 288 pixels binned to 144 pixels and where indicated sorted by heterogeneous refinement. Particles from the highest resolution reconstruction were extracted with an unbinned box size of 288 pixels and were subjected to homogeneous refinement followed by non-uniform refinement. Where indicated, particles were exported using csparc2star.py from the pyem v0.5 script package^53^, and an inclusion mask covering the 7TM domain was generated using the Segger tool in UCSF ChimeraX v1.25^54^ and the mask.py tool in pyem v0.5. Particles and mask were imported into Relion v4.0^55^ and sorted by several rounds of 3D classification without image alignment, in which the number of classes and tau factor were allowed to vary. The resulting particles were brought back into cryoSPARC and subjected to non-uniform refinement. Finally, for all datasets a local refinement using an inclusion mask covering the 7TM domain was performed. Pose/shifts Gaussian priors were used with standard deviation of rotational and shift magnitudes set as indicated in Extended data Figs 4, 5, 8, and 9.

### Site-directed mutagenesis

Generation of OR mutants was performed as previously described^56^. Forward and reverse primers coding for the mutation of interest were obtained from Integrated DNA Technologies. Two successive rounds of PCR using Phusion polymerase (F-549L, Thermo Fisher Scientific) were performed to amplify ORs with mutations. The first round of PCR generated two fragments, one containing the 5′ region upstream of the mutation site and the other containing the 3′ downstream region. The second PCR amplification joined these two fragments to produce a full open reading frame of the OR. PCR products with desired length were gel purified and cloned into the MluI and NotI sites of the mammalian expression vector pCI (Promega) that contains rho-tag. Plasmids were purified using the ZymoPure miniprep kit (D4212).

### cAMP signaling assays

The GloSensor cAMP assay (Promega) was used to determine real-time cAMP levels downstream of OR activation in HEK293T cells, as previously described^57^. HEK293T cells (authenticated by short tandem repeat profiling and tested negative for mycoplasma contamination) were cultured in minimum essential medium (MEM; Corning) supplemented by 10% FBS (Gibco), 0.5% penicillin–streptomycin (Gibco) and 0.5% amphotericin B (Gibco). Cultured HEK293T cells were plated the day before transfection at 1/10 of 100% confluence from a 100-mm plate into 96-well plates coated with poly-d-lysine (Corning) or tissue-culture coated 96-well plates with 0.001% poly-d-lysine (Sigma). For each 96-well plate, 10 μg pGloSensor-20F plasmid (Promega), 5 ug of RTP1S plasmid (only for OR1A1 and its mutants), and 75 μg of rho-tagged OR in the pCI mammalian expression vector (Promega) were transfected 18–24 h before odorant stimulation using Lipofectamine 2000 (11668019, Invitrogen) in MEM supplemented by 10% FBS. On stimulation day, plates were injected with 25 μl of GloSensor substrate (Promega) and incubated for 2 h in the dark at room temperature and in an odor-free environment. Odorants were diluted to the desired concentration in CD293 medium (Gibco) supplemented with copper (30 μM CuCl_2_; Sigma-Aldrich) and 2 mM l-glutamine (Gibco) and pH adjusted to 7.0 with a 150 mM solution of sodium hydroxide (Sigma-Aldrich). After injecting 25 μl of odorants in CD293 medium into each well, GloSensor luminescence was immediately recorded for 20 cycles of monitoring over a total period of 30 min using a BMG Labtech POLARStar Optima plate reader. The resulting luminescence activity was normalized to an empty vector negative control, and the OR response was obtained by calculating the Area Under the Curve (AUC) by summing the response from all 20 cycles. Dose-dependent responses of ORs were analyzed by fitting a least squares function to the data and by generating EC50 and efficacy using GraphPrism 10. The Area Under the dose response curve was then calculated by summing the response from each concentrations.

### Evaluating cell-surface expression

Flow cytometry was used to evaluate cell-surface expression of ORs as previously described^19^. HEK293T cells were seeded onto 35-mm plates (Greiner Bio-One) with approximately 3.5 × 105 cells (25% confluency). The cells were cultured overnight. After 18–24 h, 1,200 ng of ORs tagged with the first 20 amino acids of human rhodopsin (rho-tag) at the N-terminal ends in pCI mammalian expression vector (Promega) and 30 ng eGFP were transfected using Lipofectamine 2000 (11668019, Invitrogen). 18–24 h after transfection, the cells were detached and resuspended using Cell stripper (Corning) and then transferred into 5-ml round bottom polystyrene tubes (Falcon) on ice. The cells were spun down at 4 °C and resuspended in PBS (Gibco) containing 15 mM NaN3 (Sigma-Aldrich) and 2% FBS. (Gibco). They were stained with 1/400 (v/v) of primary antibody mouse anti-rhodopsin clone 4D2 (MABN15, Sigma-Aldrich) and allowed to incubate for 30 min, then washed with PBS containing 15 mM NaN3 and 2% FBS. The cells were spun again and then stained with 1/200 (v/v) of the phycoerythrin-conjugated donkey anti-mouse F(ab′)2 fragment antibody (715-116-150, Jackson Immunologicals) and allowed to incubate for 30 min in the dark. To label dead cells, 1/500 (v/v) of 7-amino-actinomycin D (129935, Calbiochem) was added. The cells were then immediately analyzed using a BD FACSCanto II flow cytometer with gating allowing for GFP-positive, single, spherical, viable cells, and the measured phycoerythrin fluorescence intensities were analyzed and visualized using Flowjo v10.8.1. Empty plasmid pCI is used as negative control.

### Homology model of OR1A1 and docking studies

The OR1A1 homology model was generated with the consOR1 bound to L-menthol and G protein cryo-EM structure as template using Schrödinger Maestro (version 2022-2). The consOR1 cryo-EM structure was prepared using the protein preparation wizard, which involved adding missing side chains and hydrogen atoms. Subsequently, the model was refined through hydrogen bond assignment and energy minimization. A pairwise alignment of consOR1 and OR1A1 sequences was then performed, revealing a 64% sequence identity and identifying a gap at position 194 in consOR1 compared to OR1A1. Lastly, the knowledge-based method within the build homology model module was employed to create the OR1A1 homology model, and the corresponding homology model was further energy minimized.

For docking studies of R-carvone and L-menthol into the OR1A1 homology model, we followed the Schrodnger induced fit docking protocol, with the following steps: 1. Constrained minimization of the receptor with an RMSD cutoff of 0.18 Å. 2. An RMSD alignment of the consOR1 EM structure onto the OR1A1 homology model is performed, followed by definition of a 25 Å x 25 Å x 25 Å docking grid box centered on the position of L-menthol in consOR1. This step was followed by initial Glide docking of R-carvone and L-menthol using a softened potential and removal of side chains that are within 5 Å of L-menthol. 3. A Prime side-chain prediction for each receptor-ligand pose, to rebuild the side chain conformation. 4. A Prime minimization on the receptor-ligand complex. 5. After removal of the ligand, a rigid Glide redocking is performed to re-dock the ligand back into the ligand binding site. 6. Estimation of the binding energy.

### Molecular dynamics simulations

Simulations were performed similarly to previous methods^14^ using the GROMACS package (version 2022^58^) and the CHARMM36m force field^59^. The following simulation systems were constructed: consOR1-Apo, consOR1-L-menthol bound, consOR1-L-menthol-miniGα_s_ subunit bound, consOR2-S-carvone-miniGα_s_ subunit bound, consOR2-Apo, consOR4-2MT-miniGα_s_ subunit bound, consOR4-Apo. For the miniGα_s_ subunit bound simulations, the Gβγ subunit was removed from the cryo-EM structure to reduce computational time. All ligands were parameterized by ParaChem^60^. The GPCR structures were prepared using the Maestro Schrödinger (version 2022-2) protein preparation wizard module. Missing side chains and hydrogen atoms were added, protein termini were capped with neutral acetyl and methylamide groups, and histidine states were assigned. The complex was then minimized. The simulation box was created using CHARMM-GUI^61,62^. OR1 and OR1A1 were aligned in the bilayer using the PPM 2.0 function of the Orientation of Proteins in Membranes (OPM) tool^63^, and the bilayer was filled with 75% palmitoyl-oleoyl-phosphatidylcholine (POPC) and 25% cholesteryl hemisuccinate deprotonated (CHSD). The initial positions of CHSD were taken from our previous study on OR51E2^14^. TIP3P water molecules were used for solvation, while 0.15 M potassium chloride ions were added to neutralize the system box. The final system dimensions were approximately 85 Å × 85 Å × 110 Å without the Gα subunit and 100 Å × 100 Å × 150 Å with the miniGα_s_ subunit.

The system was minimized with position restraints (10 kcal/mol/Å²) on all heavy atoms of the protein, ligand, and head group atoms of lipids, followed by a 1 ns heating step that raised the temperature from 0K to 310K in the NVT ensemble using the Nosé-Hoover thermostat. Next, a 1 μs long equilibration in the NPT ensemble was performed. During the heating step and the long equilibration, the same position restraints of 10 kcal/mol/Å² were applied for the first 1 ns, then reduced to 5 kcal/mol/Å², and gradually to 0 kcal/mol/Å² in steps of 1 kcal/mol/Å², with 5 ns of simulations per equilibration window. Afterward, a 50 ns non-restrained equilibration was conducted.

The final snapshot of the equilibration step served as the initial conformation for five production runs, which were initiated with randomly generated velocities. Pressure was coupled to a 1 bar pressure bath and controlled using the Parrinello-Rahman method^64^. Throughout all simulations, the LINCS algorithm was applied to all bonds and angles of water molecules, with a 2 fs time step employed for integration. Additionally, a 12 Å cutoff was used for non-bonded interactions, and the particle mesh Ewald method^65^ treated long-range L-J interactions. MD snapshots were saved every 20 ps, and all MD analyses were conducted on the aggregated trajectories for each system from the five runs (totaling 5 × 1000 ns = 5000 ns) using VMD (version 1.9.4), PyMOL (version 2.5), GROMACS modules (versions 2019-2022), and Python scripts.

### Ligand-receptor interactions analysis

Ligand-receptor contact frequencies were determined using the get_contacts script (https://getcontacts.github.io/). Measurements were carried out on trajectories that included solvents. All types of contacts were taken into account, encompassing water-mediated contacts as well. Contact frequencies were visualized as heatmaps using the matplotlib library.

### Ligand binding site volume calculations

To calculate the volume of the ligand-binding site, we first performed protein conformational clustering using the GROMACS cluster module. Clustering was conducted on the Cα atoms of proteins, adjusting the RMSD cutoff between 1.5 Å and 1.9 Å until the top 5-10 clusters encompassed more than 60% of all sampled points. The centroid structure of each of the top clusters was then used for ligand-binding site volume calculation. Volume calculations were executed using the Maestro SiteMap module, defining the ligand-binding pocket as being within 6 Å of the ligand. For structures obtained from Apo simulations, docking was first performed to insert L-menthol into the pocket. Subsequently, these docked structures underwent the same volume calculation protocol as the others. In the volume calculation, a more restrictive definition of hydrophobicity and fine grid was applied, and the ligand-binding site map was cropped at 4 Å from the nearest site point. The calculated volumes from the top cluster structures were utilized to compute the average and standard deviation of the ligand-binding site volume.

### Ligand flexibility analysis

Ligand RMSD values were calculated using the MDAnalysis script, which initially aligned the structure based on protein Cα atoms. Then, for each simulation frame, the RMSD matrix was computed using the coordinates of all ligand heavy atoms. Both rotational and translational movements of the ligand were taken into account. The resulting RMSD values were employed to calculate the average RMSD for ligands.

### D/E^45x51^-Y^6x55^ distance analysis

The distance between the D-Y motif was calculated using the MDAnalysis script. This distance was measured as the minimum distance between the carboxylate oxygens of D/E^45x5^^1^ (OD1, OD2 or OE1, OE2 in the CHARMM force field) and the hydroxyl oxygen (OH in the CHARMM force field) of Y^6x55^ in consOR1, consOR2 and consOR4 production trajectories, respectively. The time evaluation of distance from a selected velocity was plotted as a moving average and rolling standard deviation using the matplotlib and scipy library. The overall distances from the production trajectory was represented as a violin plot using matplotlib.

### Phylogenetic tree, sequence identity and structure comparison

On R 4.3.1, alignment reading and matrix of distance between sequences (by sequence identity) calculation were performed with the Biostrings^66^ and seqinr^67^ packages. Neighbor-Joining tree and tree visualization were realized with packages ape^68^ and ggtree^69^ and the tree is plotted unrooted with the daylight method. Sequence identity and RMSD between structures were calculated with the package bio3D and graphs were made with pheatmap and gtools packages. Conserved positions in aligned sequences of Class I and Class II ORs were visualized with WebLogo3^70^.

## Data Availability

Coordinates for consOR51, consOR1, consOR2, and consOR4 have been deposited in the RCSB PDB under accession codes 8UXV, 8UXY, 8UY0, and 8UYQ, respectively. EM density maps for consOR51, consOR1, consOR2, and consOR4 have been deposited in the Electron Microscopy Data Bank under accession codes EMD-42786, EMD-42789, EMD-42791, and EMD-42817 respectively. The MD simulation trajectories have been deposited in the GPCRmd database under access codes XXXX.

## Acknowledgements

This work was supported by the National Institutes of Health (NIH) grant R01DC020353 (H.M., N.V., and A.M.) and K99DC018333 (C.A.D.M.). Cryo-EM equipment at UCSF is partially supported by NIH grants S10OD020054 and S10OD021741. Some of this work was performed at the Stanford-SLAC Cryo-EM Center (S2C2), which is supported by the National Institutes of Health Common Fund Transformative High-Resolution Cryo-Electron Microscopy program (U24 GM129541). This project was funded by the UCSF Program for Breakthrough Biomedical Research, funded in part by the Sandler Foundation. A.M. acknowledges support from the Edward Mallinckrodt, Jr. Foundation and the Vallee Foundation. A.M. is a Chan Zuckerberg Biohub San Francisco Investigator.

## Contributions

C.A.D.M. and H.M. designed the consensus OR strategy, and with A.M., outlined a structure determination strategy for consensus ORs. The study was also designed by C.B.B., N.M., W.J.C.v.d.V, and N.V. Consensus constructs were designed and cloned by C.A.D.M. with input from A.M. C.A.D.M generated the phylogenetic trees. C.B.B. led the effort for structure determination, including cloning constructs, preparing baculoviruses, expressing and purifying G protein complexing reagents. With. C.L.D.T., C.B.B. established conditions to biochemically purify and stabilize consOR1, consOR2, and consOR4 complexes, identified optimal cryo-EM grid preparation procedures, and collected cryo-EM datasets for structure determination. L.L. assisted with protein purifications. A.M. purified the consOR51 complex and collected cryo-EM data with help from B.F. C.B.B. determined high-resolution cryo-EM maps by extensive image processing with input from A.M. and C.L.D.T. A.M. and C.B.B. built and refined models of consOR complexes. C.A.D.M. and J.T. analyzed OR models and sequences to design and clone OR mutants. C.A.D.M., J.T., and I.O. performed Glosensor signaling experiments for OR functional activity, and C.A.D.M., J.T., I.T and I.O. generated OR cell surface expression data by flow cytometry with input from H.M. C.A.D.M and J.T. analyzed and prepared figures and tables for signaling and flow cytometry data. N.M. set up molecular dynamics simulations, ligand docking, and performed binding pocket volume calculations. W.J.C.v.d.V created the homology model of OR1A1. N.M. and W.J.C.v.d.V. analyzed simulation trajectories and prepared figures describing simulation data. N.M., W.J.C.v.d.V. and N.V. provided mechanistic insight from simulation data. C.A.D.M. performed the comparative structure analysis. C.A.D.M., N.M., and A.M. wrote an initial draft of the manuscript and generated figures with contributions from all authors. Further edits to the manuscript were provided by W.J.C.v.d.V., N.M., N.V., and H.M. The overall project was supervised and funded by C.A.D.M., N.V., H.M., and A.M.

## REFERENCES

1. Malnic, B., Godfrey, P. A. & Buck, L. B. The human olfactory receptor gene family. Proc. Natl. Acad. Sci. U. S. A. 101, 2584–2589 (2004).

2. Bjarnadóttir, T. K. et al. Comprehensive repertoire and phylogenetic analysis of the G protein-coupled receptors in human and mouse. Genomics 88, 263–273 (2006).

3. Glusman, G., Yanai, I., Rubin, I. & Lancet, D. The complete human olfactory subgenome. Genome Res. 11, 685–702 (2001).

4. Buck, L. & Axel, R. A novel multigene family may encode odorant receptors: a molecular basis for odor recognition. Cell 65, 175–187 (1991).

5. Liberles, S. D. & Buck, L. B. A second class of chemosensory receptors in the olfactory epithelium. Nature 442, 645–650 (2006).

6. Olender, T., Jones, T. E. M., Bruford, E. & Lancet, D. A unified nomenclature for vertebrate olfactory receptors. BMC Evol. Biol. 20, 42 (2020).

7. Malnic, B., Hirono, J., Sato, T. & Buck, L. B. Combinatorial receptor codes for odors. Cell 96, 713–723 (1999).

8. Saito, H., Chi, Q., Zhuang, H., Matsunami, H. & Mainland, J. D. Odor coding by a Mammalian receptor repertoire. Sci. Signal. 2, ra9 (2009).

9. Cichy, A., Shah, A., Dewan, A., Kaye, S. & Bozza, T. Genetic Depletion of Class I Odorant Receptors Impacts Perception of Carboxylic Acids. Curr. Biol. 29, 2687–2697.e4 (2019).

10. Dewan, A., Pacifico, R., Zhan, R., Rinberg, D. & Bozza, T. Non-redundant coding of aversive odours in the main olfactory pathway. Nature 497, 486–489 (2013).

11. Niimura, Y. On the origin and evolution of vertebrate olfactory receptor genes: comparative genome analysis among 23 chordate species. Genome Biol. Evol. 1, 34–44 (2009).

12. Bear, D. M., Lassance, J.-M., Hoekstra, H. E. & Datta, S. R. The Evolving Neural and Genetic Architecture of Vertebrate Olfaction. Curr. Biol. 26, R1039–R1049 (2016).

13. Freitag, J., Krieger, J., Strotmann, J. & Breer, H. Two classes of olfactory receptors in Xenopus laevis. Neuron 15, 1383–1392 (1995).

14. Billesbølle, C. B. et al. Structural basis of odorant recognition by a human odorant receptor. Nature 615, 742–749 (2023).

15. 15. Gusach, A., et al. Molecular recognition of an aversive odorant by the murine trace amine-associated receptor TAAR7f. *bioRxiv* (2023) doi:10.1101/2023.07.07.547762.

16. Guo, L. et al. Structural basis of amine odorant perception by a mammal olfactory receptor. Nature 618, 193–200 (2023).

17. Lu, M., Echeverri, F. & Moyer, B. D. Endoplasmic reticulum retention, degradation, and aggregation of olfactory G-protein coupled receptors. Traffic 4, 416–433 (2003).

18. Saito, H., Kubota, M., Roberts, R. W., Chi, Q. & Matsunami, H. RTP family members induce functional expression of mammalian odorant receptors. Cell 119, 679–691 (2004).

19. Zhuang, H. & Matsunami, H. Evaluating cell-surface expression and measuring activation of mammalian odorant receptors in heterologous cells. Nat. Protoc. 3, 1402–1413 (2008).

20. Noe, F. et al. IL-6-HaloTag® enables live-cell plasma membrane staining, flow cytometry, functional expression, and de-orphaning of recombinant odorant receptors. J Biol Methods 4, e81 (2017).

21. Sternke, M., Tripp, K. W. & Barrick, D. Consensus sequence design as a general strategy to create hyperstable, biologically active proteins. Proc. Natl. Acad. Sci. U. S. A. 116, 11275–11284 (2019).

22. Desjarlais, J. R. & Berg, J. M. Use of a zinc-finger consensus sequence framework and specificity rules to design specific DNA binding proteins. Proc. Natl. Acad. Sci. U. S. A. 90, 2256–2260 (1993).

23. Porebski, B. T. & Buckle, A. M. Consensus protein design. Protein Eng. Des. Sel. 29, 245– 251 (2016).

24. Steipe, B., Schiller, B., Plückthun, A. & Steinbacher, S. Sequence statistics reliably predict stabilizing mutations in a protein domain. J. Mol. Biol. 240, 188–192 (1994).

25. Lehmann, M. et al. From DNA sequence to improved functionality: using protein sequence comparisons to rapidly design a thermostable consensus phytase. Protein Eng. 13, 49–57 (2000).

26. Ikegami, K. et al. Structural instability and divergence from conserved residues underlie intracellular retention of mammalian odorant receptors. Proc. Natl. Acad. Sci. U. S. A. 117, 2957–2967 (2020).

27. Nehmé, R. et al. Mini-G proteins: Novel tools for studying GPCRs in their active conformation. PLoS One 12, e0175642 (2017).

28. Ballesteros, J. A. & Weinstein, H. [19] Integrated methods for the construction of three-dimensional models and computational probing of structure-function relations in G protein-coupled receptors. in Methods in Neurosciences (ed. Sealfon, S. C.) vol. 25 366–428 (Academic Press, 1995).

29. de March, C. A., Kim, S.-K., Antonczak, S., Goddard, W. A., 3rd & Golebiowski, J. G protein-coupled odorant receptors: From sequence to structure. Protein Sci. 24, 1543–1548 (2015).

30. Isberg, V. et al. Generic GPCR residue numbers - aligning topology maps while minding the gaps. Trends Pharmacol. Sci. 36, 22–31 (2015).

31. de March, C. A. et al. Conserved Residues Control Activation of Mammalian G Protein-Coupled Odorant Receptors. J. Am. Chem. Soc. 137, 8611–8616 (2015).

32. Pluznick, J. L. et al. Olfactory receptor responding to gut microbiota-derived signals plays a role in renin secretion and blood pressure regulation. Proc. Natl. Acad. Sci. U. S. A. 110, 4410–4415 (2013).

33. Shayya, H. J. et al. ER stress transforms random olfactory receptor choice into axon targeting precision. Cell 185, 3896–3912.e22 (2022).

34. Mainland, J. D., Li, Y. R., Zhou, T., Liu, W. L. L. & Matsunami, H. Human olfactory receptor responses to odorants. Sci Data 2, 150002 (2015).

35. Kajiya, K. et al. Molecular bases of odor discrimination: Reconstitution of olfactory receptors that recognize overlapping sets of odorants. J. Neurosci. 21, 6018–6025 (2001).

36. Grosmaitre, X. et al. SR1, a mouse odorant receptor with an unusually broad response profile. J. Neurosci. 29, 14545–14552 (2009).

37. Schmiedeberg, K. et al. Structural determinants of odorant recognition by the human olfactory receptors OR1A1 and OR1A2. J. Struct. Biol. 159, 400–412 (2007).

38. Ma, N., Lee, S. & Vaidehi, N. Activation Microswitches in Adenosine Receptor A2A Function as Rheostats in the Cell Membrane. Biochemistry 59, 4059–4071 (2020).

39. Dror, R. O. et al. Activation mechanism of the *β*_2_-adrenergic receptor. Proc. Natl. Acad. Sci. U. S. A. 108, 18684–18689 (2011).

40. Lee, S., Nivedha, A. K., Tate, C. G. & Vaidehi, N. Dynamic Role of the G Protein in Stabilizing the Active State of the Adenosine A2A Receptor. Structure 27, 703–712.e3 (2019).

41. Li, Q. et al. Non-classical amine recognition evolved in a large clade of olfactory receptors. Elife 4, e10441 (2015).

42. Del Mármol, J., Yedlin, M. A. & Ruta, V. The structural basis of odorant recognition in insect olfactory receptors. Nature 597, 126–131 (2021).

43. Butterwick, J. A. et al. Cryo-EM structure of the insect olfactory receptor Orco. Nature 560, 447–452 (2018).

44. Jumper, J. et al. Highly accurate protein structure prediction with AlphaFold. Nature 596, 583–589 (2021).

45. Bender, B. J., Marlow, B. & Meiler, J. Improving homology modeling from low-sequence identity templates in Rosetta: A case study in GPCRs. PLoS Comput. Biol. 16, e1007597 (2020).

46. Rutherford, S. L. & Lindquist, S. Hsp90 as a capacitor for morphological evolution. Nature 396, 336–342 (1998).

47. Wyganowski, K. T., Kaltenbach, M. & Tokuriki, N. GroEL/ES buffering and compensatory mutations promote protein evolution by stabilizing folding intermediates. J. Mol. Biol. 425, 3403–3414 (2013).

48. Agozzino, L. & Dill, K. A. Protein evolution speed depends on its stability and abundance and on chaperone concentrations. Proc. Natl. Acad. Sci. U. S. A. 115, 9092–9097 (2018).

49. Faust, B. et al. Autoantibody and hormone activation of the thyrotropin G protein-coupled receptor. bioRxiv 2022.01.06.475289 (2022) doi:10.1101/2022.01.06.475289.

50. Mastronarde, D. N. SerialEM: A Program for Automated Tilt Series Acquisition on Tecnai Microscopes Using Prediction of Specimen Position. Microsc. Microanal. 9, 1182–1183 (2003).

51. Zheng, S. Q. et al. MotionCor2: anisotropic correction of beam-induced motion for improved cryo-electron microscopy. Nat. Methods 14, 331–332 (2017).

52. Punjani, A., Rubinstein, J. L., Fleet, D. J. & Brubaker, M. A. cryoSPARC: algorithms for rapid unsupervised cryo-EM structure determination. Nat. Methods 14, 290–296 (2017).

53. Asarnow, D., Palovcak, E. & Cheng, Y. asarnow/pyem: UCSF pyem v0.5. (2019). doi:10.5281/zenodo.3576630.

54. Pettersen, E. F. et al. UCSF ChimeraX: Structure visualization for researchers, educators, and developers. Protein Sci. 30, 70–82 (2021).

55. Scheres, S. H. W. RELION: implementation of a Bayesian approach to cryo-EM structure determination. J. Struct. Biol. 180, 519–530 (2012).

56. Bushdid, C., de March, C. A., Matsunami, H. & Golebiowski, J. Numerical Models and In Vitro Assays to Study Odorant Receptors. Methods Mol. Biol. 1820, 77–93 (2018).

57. Zhang, Y., Pan, Y., Matsunami, H. & Zhuang, H. Live-cell Measurement of Odorant Receptor Activation Using a Real-time cAMP Assay. J. Vis. Exp. (2017) doi:10.3791/55831.

58. Berendsen, H. J. C., van der Spoel, D. & van Drunen, R. GROMACS: A message-passing parallel molecular dynamics implementation. Comput. Phys. Commun. 91, 43–56 (1995).

59. Huang, J. et al. CHARMM36m: an improved force field for folded and intrinsically disordered proteins. Nat. Methods 14, 71–73 (2017).

60. Vanommeslaeghe, K. et al. CHARMM general force field: A force field for drug-like molecules compatible with the CHARMM all-atom additive biological force fields. J. Comput. Chem. 31, 671–690 (2010).

61. Jo, S., Kim, T., Iyer, V. G. & Im, W. CHARMM-GUI: a web-based graphical user interface for CHARMM. J. Comput. Chem. 29, 1859–1865 (2008).

62. Jo, S., Lim, J. B., Klauda, J. B. & Im, W. CHARMM-GUI Membrane Builder for mixed bilayers and its application to yeast membranes. Biophys. J. 97, 50–58 (2009).

63. Lomize, M. A., Pogozheva, I. D., Joo, H., Mosberg, H. I. & Lomize, A. L. OPM database and PPM web server: resources for positioning of proteins in membranes. Nucleic Acids Res. 40, D370–6 (2012).

64. Parrinello, M. & Rahman, A. Polymorphic transitions in single crystals: A new molecular dynamics method. J. Appl. Phys. 52, 7182–7190 (1981).

65. Darden, T., York, D. & Pedersen, L. Particle mesh Ewald: An *N*⋅log(*N*) method for Ewald sums in large systems. J. Chem. Phys. 98, 10089–10092 (1993).

66. Pagès, H., Aboyoun, P., Gentleman, R. & DebRoy, S. Biostrings: Efficient manipulation of biological 370 strings. 10.18129/B9. bioc. Preprint at (2022).

67. Charif, D. & Lobry, J. R. SeqinR 1.0-2: A Contributed Package to the R Project for Statistical Computing Devoted to Biological Sequences Retrieval and Analysis. in Structural Approaches to Sequence Evolution: Molecules, Networks, Populations (eds. Bastolla, U., Porto, M., Roman, H. E. & Vendruscolo, M.) 207–232 (Springer Berlin Heidelberg, 2007).

68. Paradis, E. & Schliep, K. ape 5.0: an environment for modern phylogenetics and evolutionary analyses in R. Bioinformatics 35, 526–528 (2019).

69. Xu, S., et al. *Ggtree* : A serialized data object for visualization of a phylogenetic tree and annotation data. iMeta (2022) doi:10.1002/imt2.56.

70. Crooks, G. E., Hon, G., Chandonia, J.-M. & Brenner, S. E. WebLogo: a sequence logo generator. Genome Res. 14, 1188–1190 (2004).

71. Dang, S. et al. Cryo-EM structures of the TMEM16A calcium-activated chloride channel. Nature 552, 426–429 (2017).

