## Supplementary Information for "Engineered odorant receptors illuminate structural principles of odor discrimination"

**SUPPLEMENTARY FIGURES:**

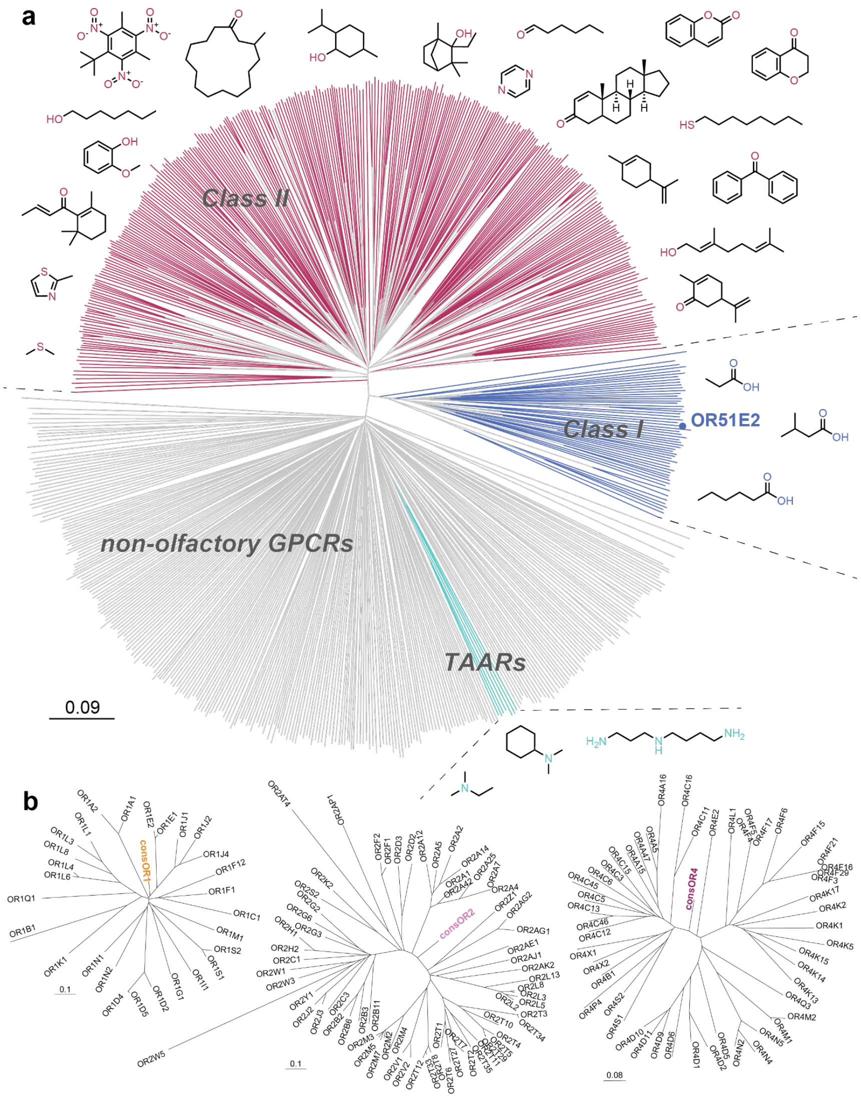


**Supplementary Figure 1. Phylogenetic tree of human Class A GPCRs and consORs. a)** Olfactory GPCRs are highlighted in pink (Class II ORs), blue (Class I ORs) and cyan (TAARs). Non-olfactory GPCRs are represented with gray lines. A subset of ligands recognized by the olfactory GPCRs are presented, with heteroatoms colored by the olfactory GPCR families responsible for detection. OR51E2, the only human odorant receptor known structure is highlighted. On the bottom left, the scale represents the amount of amino acid change for a set distance. **b)** Phylogenetic trees for human OR subfamilies OR1, OR2 and OR4. Consensus ORs (consORs) are indicated in color.

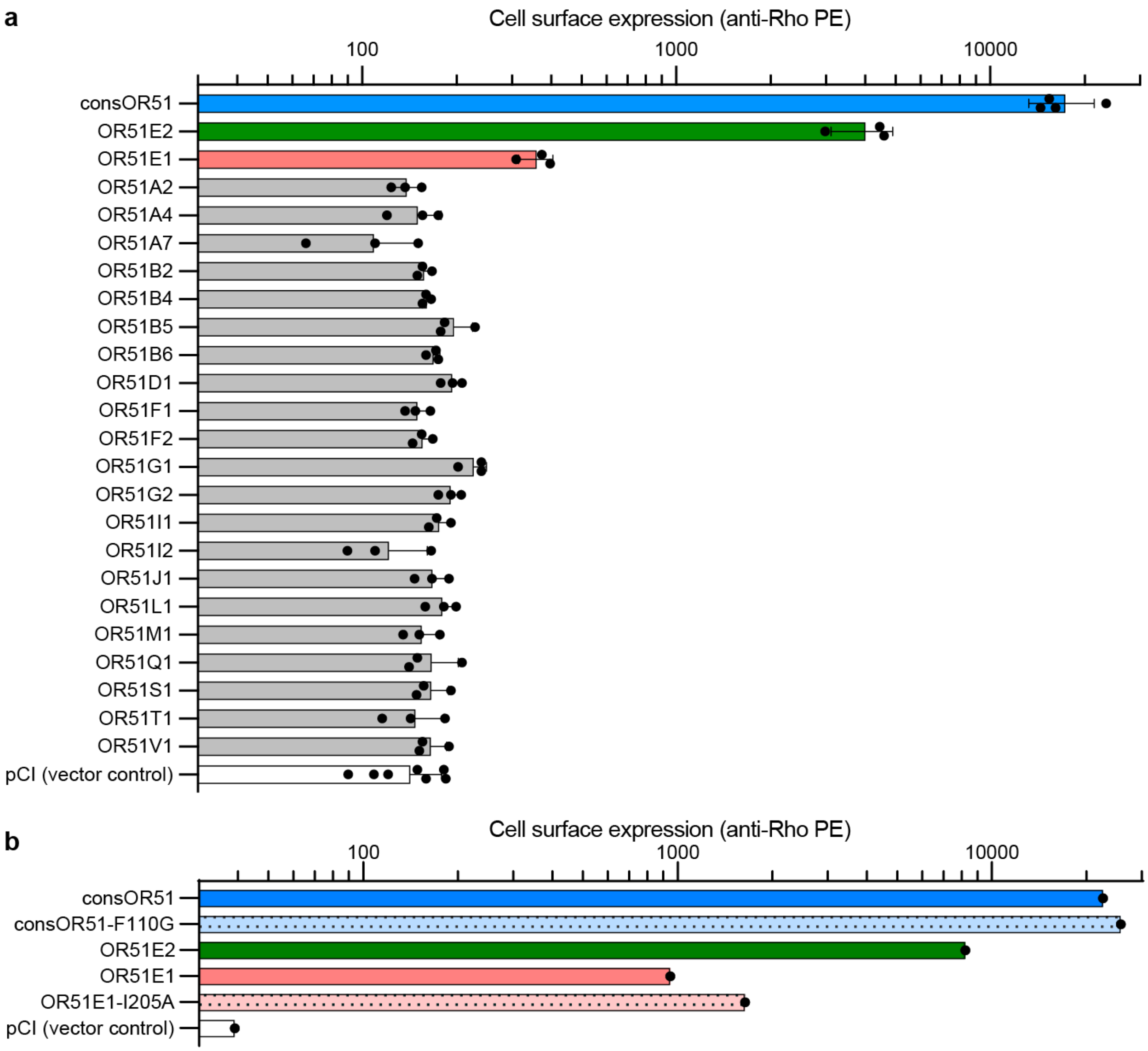


**Supplementary Figure 2. Expression of OR51 family members. a)** Cell surface expression of OR51 family members monitored by flow cytometry. **b)** Cell surface expression of consOR51 and F110G mutant as well as OR51E1 and its mutant I205A monitored by flow cytometry.

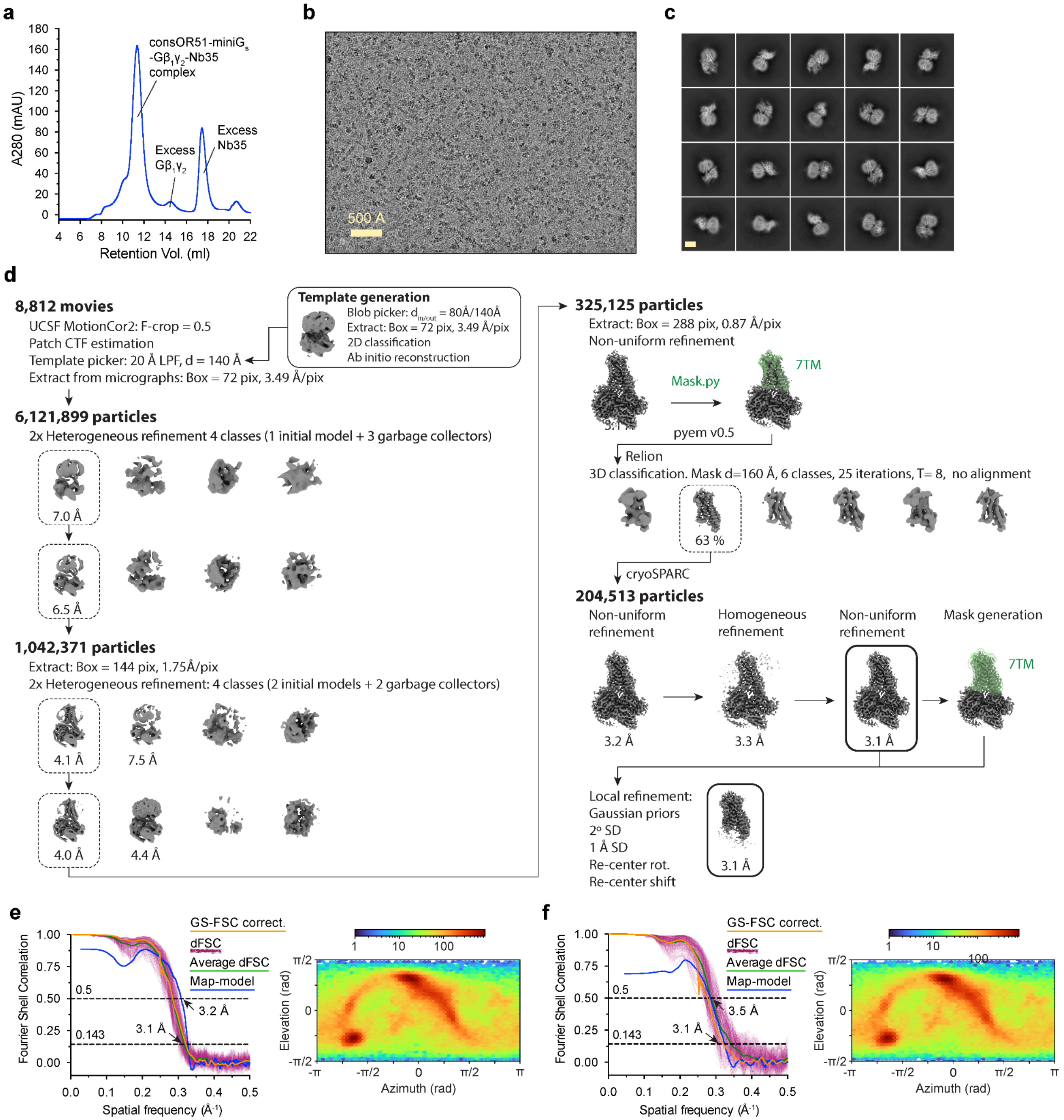


**Supplementary Figure 3. Cryo-EM data processing for consOR51-$G_s$. a)** Size exclusion chromatogram of consOR1-$G_s$ sample. **b)** A representative cryo-EM micrograph from the curated consOR51-$G_s$ dataset (n = 8,812) obtained from a Titan Krios microscope. **c)** A subset of highly populated, reference-free 2D-class averages are shown. Scale bar is 50 Å. **d)** Schematic showing the image processing workflow for consOR51-$G_s$. Initial processing was performed using UCSF MotionCor2 and cryoSPARC. Particles were then transferred using the pyem script package[53] to RELION for alignment-free 3D classification. Finally, particles were processed in cryoSPARC using the non-uniform and local refinement tools. Dashed boxes indicate selected classes, and 3D volumes of classes and refinements are shown along with global Gold-standard Fourier Shell Coefficient (GSFSC) resolutions. **e, f)** Map validation for the consOR51-$G_s$ (**e**) globally refined, and (**f**) locally refined cryo-EM maps. Gold-standard Fourier shell correlation (FSC) curves are calculated in cryoSPARC, and shown together with directional FSC (dFSC) curves generated with dfsc.0.0.1.py as previously described[71]. Map-model correlations calculated in the Phenix suite are also shown. Arrows indicate map and map-model resolution estimates at 0.143 and 0.5 correlation respectively. Euler angle distributions calculated in cryoSPARC are also provided for each map.

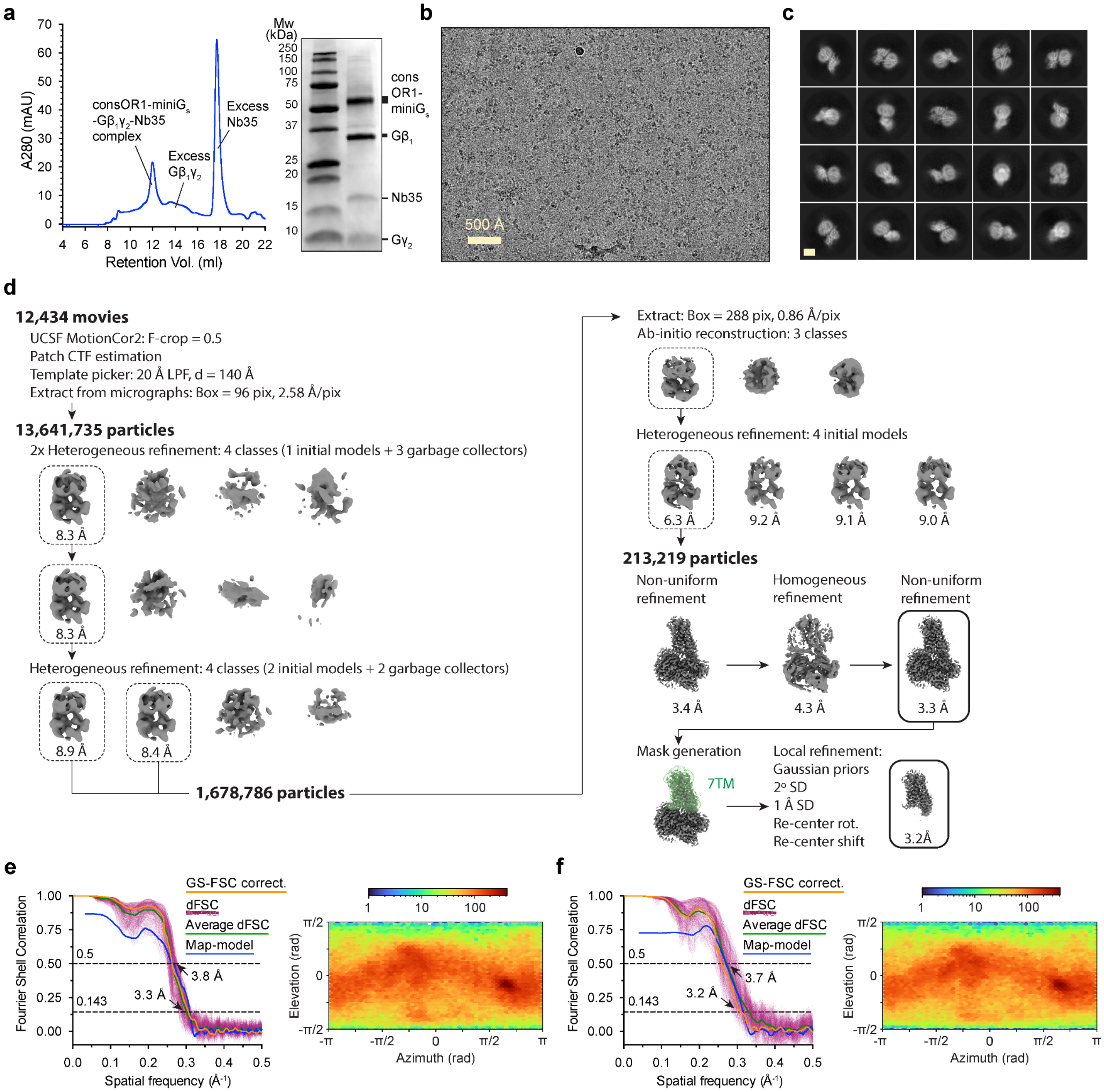


**Supplementary Figure 4. Cryo-EM data processing for consOR1-$G_s$. a)** Size exclusion chromatogram and SDS-PAGE of consOR1-$G_s$ sample. **b)** A representative cryo-EM micrograph from the curated consOR1-$G_s$ dataset (n = 12,434) obtained from a Titan Krios microscope. **c)** A subset of highly populated, reference-free 2D-class averages are shown. Scale bar is 50 Å. **d)** Schematic showing the image processing workflow for consOR1-$G_s$. Initial processing was performed using UCSF MotionCor2 and cryoSPARC. Particles were then transferred using the pyem script package[53] to RELION for alignment-free 3D classification. Finally, particles were processed in cryoSPARC using the non-uniform and local refinement tools. Dashed boxes indicate selected classes, and 3D volumes of classes and refinements are shown along with global Gold-standard Fourier Shell Coefficient (GSFSC) resolutions. **e, f)** Map validation for the consOR1-$G_s$ (**e**) globally refined, and (**f**) locally refined cryo-EM maps. Gold-standard Fourier shell correlation (FSC) curves are calculated in cryoSPARC, and shown together with directional FSC (dFSC) curves generated with dfsc.0.0.1.py as previously described[71]. Map-model correlations calculated in the Phenix suite are also shown. Arrows indicate map and map-model resolution estimates at 0.143 and 0.5 correlation respectively. Euler angle distributions calculated in cryoSPARC are also provided for each map.

**a**

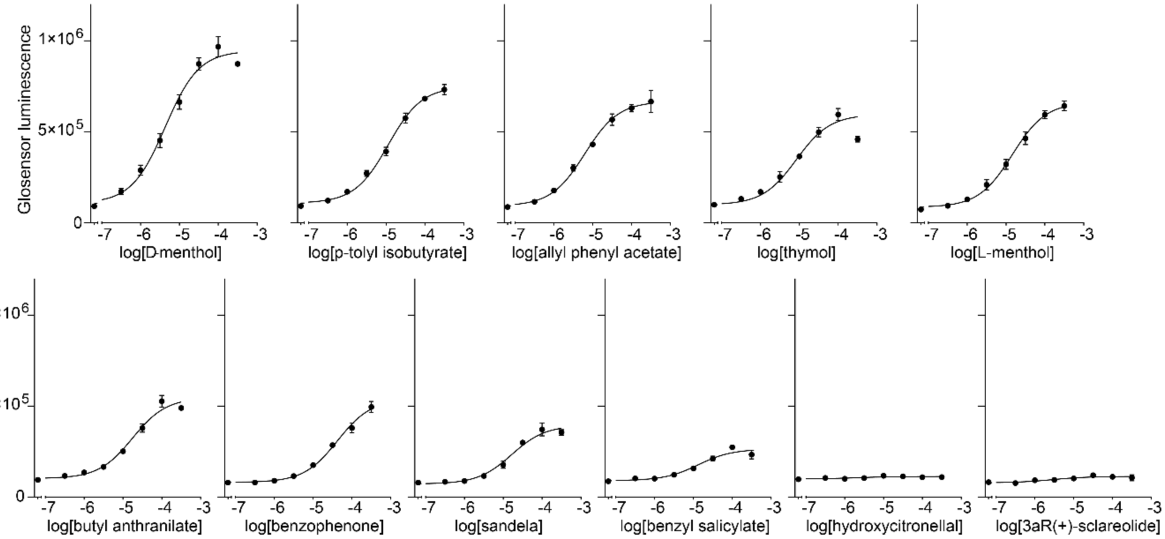


**b**

| consOR1 | Bottom | | Top | | LogEC50 | | EC50 | Span | | $R^2$ fit | AUC | |
|---|---|---|---|---|---|---|---|---|---|---|---|---|
| | Mean | SEM | Mean | SEM | Mean | SEM | Mean | Mean | SEM | | Mean | SEM |
| D-menthol | 107010 | 20141 | 945375 | 18850 | -5.37 | 0.06 | 4.31E-06 | 838364 | 24928 | 0.98 | 6.36 | 0.15 |
| 3aR(+)-sclareolide | non fit | non fit | non fit | non fit | non fit | non fit | non fit | non fit | non fit | 0.64 | 1.14 | 0.05 |
| L-menthol | 85119 | 8174 | 662189 | 12490 | -4.84 | 0.04 | 1.45E-05 | 577070 | 13406 | 0.99 | 3.66 | 0.11 |
| allyl phenyl acetate | 91593 | 10225 | 666784 | 11017 | -5.20 | 0.05 | 6.26E-06 | 575192 | 13511 | 0.99 | 4.32 | 0.09 |
| benzophenone | 80746 | 5309 | 537852 | 14300 | -4.37 | 0.05 | 4.24E-05 | 457107 | 13988 | 0.99 | 2.47 | 0.06 |
| benzyl salicylate | 90051 | 6715 | 263641 | 9864 | -4.88 | 0.12 | 1.32E-05 | 173590 | 10710 | 0.93 | 1.87 | 0.06 |
| butyl anthranilate | 101550 | 9232 | 545631 | 15440 | -4.76 | 0.07 | 1.76E-05 | 444081 | 16187 | 0.98 | 3.13 | 0.04 |
| hydroxycitronellal | non fit | non fit | non fit | non fit | non fit | non fit | non fit | non fit | non fit | 0.30 | 1.24 | 0.03 |
| p-tolyl isobutyrate | 107472 | 7739 | 748344 | 10631 | -4.94 | 0.04 | 1.14E-05 | 640873 | 11795 | 0.99 | 4.41 | 0.09 |
| sandela | 71822 | 7483 | 394389 | 12132 | -4.78 | 0.07 | 1.64E-05 | 322567 | 12817 | 0.97 | 2.28 | 0.07 |
| thymol | 101177 | 18881 | 543827 | 19649 | -5.24 | 0.11 | 5.72E-06 | 442650 | 24520 | 0.94 | 3.73 | 0.04 |
| no odor | non fit | non fit | non fit | non fit | non fit | non fit | non fit | non fit | non fit | 0.56 | 1.00 | 0.01 |

**c**

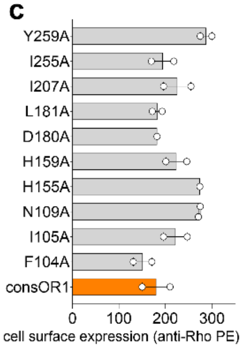


**d**

| L-menthol | Bottom | | Top | | LogEC50 | | EC50 | Span | | $R^2$ fit | AUC | |
|---|---|---|---|---|---|---|---|---|---|---|---|---|
| | Mean | SEM | Mean | SEM | Mean | SEM | Mean | Mean | SEM | | Mean | SEM |
| OR1A1 | 1.91 | 0.29 | 19.18 | 0.39 | -4.79 | 0.05 | 1.62E-05 | 17.27 | 0.43 | 0.99 | 1.00 | 0.02 |
| OR1A1 M104A | 5.55 | 0.23 | 16.79 | 0.45 | -4.47 | 0.07 | 3.36E-05 | 11.25 | 0.45 | 0.98 | 0.99 | 0.02 |
| OR1A1 I105A | 1.14 | 0.23 | 11.16 | 0.27 | -4.60 | 0.06 | 2.50E-05 | 10.03 | 0.32 | 0.98 | 0.59 | 0.01 |
| OR1A1 N109A | no fit | no fit | no fit | no fit | no fit | no fit | no fit | no fit | no fit | 0.73 | 0.12 | 0.00 |
| OR1A1 N155A | 2.44 | 0.40 | 12.90 | 0.42 | -4.72 | 0.10 | 1.90E-05 | 10.46 | 0.52 | 0.95 | 0.78 | 0.01 |
| OR1A1 H159A | 1.31 | 0.15 | 7.26 | 0.29 | -4.11 | 0.09 | 7.79E-05 | 5.95 | 0.29 | 0.96 | 0.34 | 0.01 |
| OR1A1 I181A | 1.12 | 0.07 | 6.86 | 0.12 | -4.30 | 0.04 | 4.98E-05 | 5.74 | 0.12 | 0.99 | 0.34 | 0.01 |
| OR1A1 M198A | 1.83 | 0.20 | 12.80 | 0.17 | -5.00 | 0.04 | 9.97E-06 | 10.97 | 0.24 | 0.99 | 0.82 | 0.02 |
| OR1A1 I205A | 0.84 | 0.05 | 6.04 | 0.16 | -3.83 | 0.05 | 1.47E-04 | 5.20 | 0.15 | 0.99 | 0.24 | 0.00 |
| OR1A1 F206A | no fit | no fit | no fit | no fit | no fit | no fit | no fit | no fit | no fit | 0.00 | 0.09 | 0.00 |
| OR1A1 V254A | 0.94 | 0.08 | 4.32 | 0.10 | -4.47 | 0.07 | 3.36E-05 | 3.38 | 0.11 | 0.98 | 0.25 | 0.01 |
| OR1A1 Y258A | 0.97 | 0.19 | 6.31 | 0.21 | -4.68 | 0.09 | 2.09E-05 | 5.35 | 0.25 | 0.96 | 0.36 | 0.02 |

| R-carvone | Bottom | | Top | | LogEC50 | | EC50 | Span | | $R^2$ fit | AUC | |
|---|---|---|---|---|---|---|---|---|---|---|---|---|
| | Mean | SEM | Mean | SEM | Mean | SEM | Mean | Mean | SEM | | Mean | SEM |
| OR1A1 | 2.749 | 0.468 | 21.71 | 0.309 | -6.82 | 0.049 | 1.52E-07 | 18.96 | 0.522 | 0.985 | 1.00 | 0.02 |
| OR1A1 M104A | 6.039 | 0.25 | 20.17 | 0.378 | -5.85 | 0.055 | 1.41E-06 | 14.13 | 0.407 | 0.985 | 0.80 | 0.01 |
| OR1A1 I105A | 1.755 | 0.035 | 10.59 | 0.838 | -4.45 | 0.072 | 3.58E-05 | 8.836 | 0.825 | 0.992 | 0.18 | 0.00 |
| OR1A1 N109A | 1.006 | 0.075 | 12.65 | 0.399 | -4.98 | 0.04 | 1.05E-05 | 11.65 | 0.384 | 0.994 | 0.23 | 0.00 |
| OR1A1 N155A | 2.564 | 0.18 | 21.33 | 0.472 | -5.39 | 0.04 | 4.09E-06 | 18.76 | 0.463 | 0.992 | 0.55 | 0.01 |
| OR1A1 H159A | 2.043 | 0.08 | 20.55 | 0.35 | -5.08 | 0.024 | 8.31E-06 | 18.51 | 0.337 | 0.998 | 0.42 | 0.01 |
| OR1A1 I181A | 0.768 | 0.051 | 8.737 | 0.28 | -4.96 | 0.041 | 1.09E-05 | 7.969 | 0.269 | 0.994 | 0.16 | 0.01 |
| OR1A1 M198A | 1.916 | 0.467 | 22.2 | 0.354 | -6.63 | 0.049 | 2.36E-07 | 20.28 | 0.538 | 0.986 | 0.94 | 0.01 |
| OR1A1 I205A | 1.693 | 0.318 | 28.3 | 1.411 | -5.07 | 0.067 | 8.49E-06 | 26.61 | 1.358 | 0.982 | 0.51 | 0.02 |
| OR1A1 F206A | non fit | non fit | non fit | non fit | non fit | non fit | non fit | non fit | non fit | 0.923 | 0.08 | 0.01 |
| OR1A1 V254A | 1.507 | 0.148 | 24 | 0.92 | -4.91 | 0.046 | 1.23E-05 | 22.5 | 0.888 | 0.993 | 0.39 | 0.01 |
| OR1A1 Y258A | 0.899 | 0.064 | 10.27 | 0.619 | -4.73 | 0.064 | 1.85E-05 | 9.369 | 0.602 | 0.989 | 0.16 | 0.00 |

**e**

| | Δ Fold change *AUC(R-carvone) - AUC(L-menthol)* | |
|---|---|---|
| | mean | SEM |
| OR1A1 | 0.00 | 0.02 |
| OR1A1 M104A | -0.19 | 0.02 |
| OR1A1 I105A | -0.41 | 0.01 |
| OR1A1 N109A | 0.11 | 0.00 |
| OR1A1 N155A | -0.23 | 0.01 |
| OR1A1 H159A | 0.07 | 0.01 |
| OR1A1 I181A | -0.19 | 0.01 |
| OR1A1 M198A | 0.12 | 0.02 |
| OR1A1 I205A | 0.28 | 0.02 |
| OR1A1 F206A | -0.01 | 0.00 |
| OR1A1 V254A | 0.14 | 0.01 |
| OR1A1 Y258A | -0.21 | 0.01 |

**f**

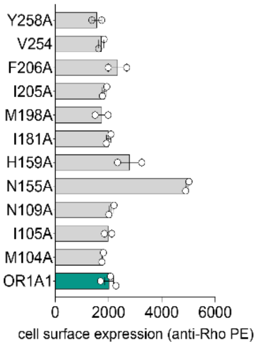


875

**Supplementary Figure 5. OR1 and OR1A1 mutants activity against odorants. a)** consOR1 response in cAMP accumulation assay to 11 odorant stimulations in dose response (concentration in mol.$L^{-1}$). **b)** Table of AUC normalized to no odor, potency and efficacy for consOR1 against the 11 odorants. **c)** Cell surface expression of consOR1 and its mutants evaluated in flow cytometry. **d)** Table of AUC normalized to the *wt*, potency and efficacy for OR1A1 and its mutants against L-menthol and R-carvone. **e)** Table of Delta Area Under Curve between R-carvone and L-menthol for OR1A1 and its mutants. **f)** Cell surface expression of OR1A1 and its mutants evaluated in flow cytometry.

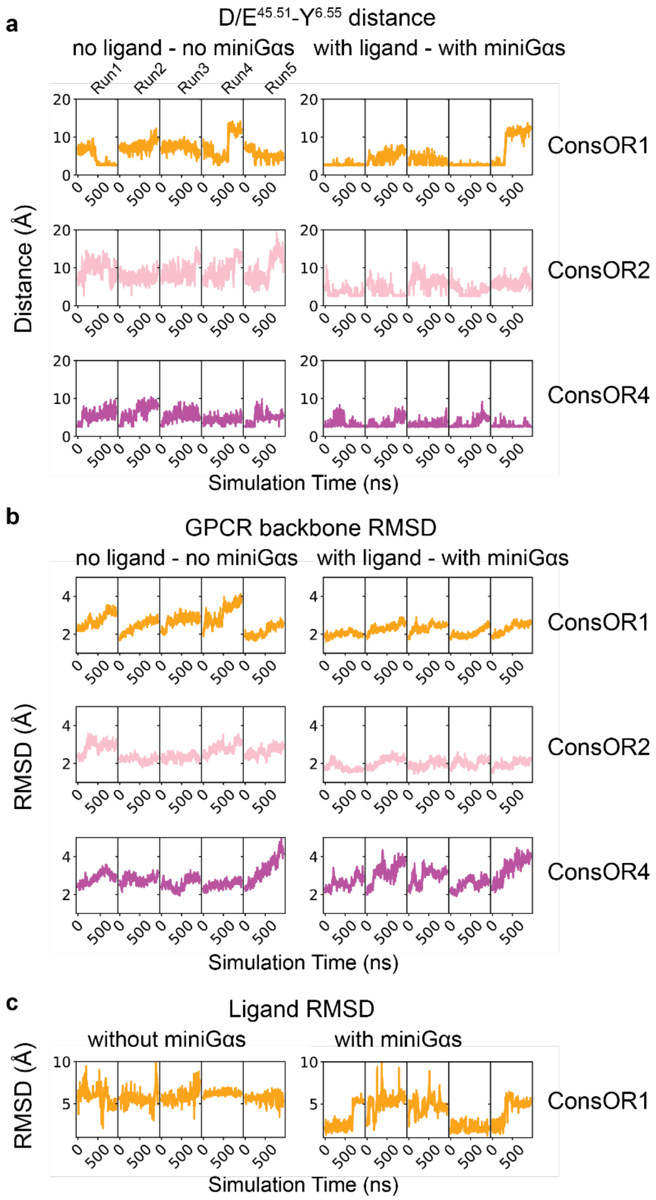


**Supplementary Figure 6. Time evolution of D/E$^{45x51}$-Y$^{6x55}$ distance and GPCR/Ligand RMSD from MD simulations .** All measurements were performed on MD trajectories by skipping every 100 frames. The raw data points were plotted transparent. A fitted curve was plotted as a solid line by using a smoothing window of 50 sampling points. **a)** The distance between D/E$^{45x51}$ and Y$^{6x55}$ from MD simulations without ligand/miniGαs and with ligand/miniGαs states for consOR1, consOR2, consOR4. The atoms involved in this measurement are described in the method section. **b)** The GPCR backbone atoms RMSD from MD simulations without ligand/miniGαs and with ligand/miniGαs states for consOR1, consOR2, consOR4. **c)**

The ligand RMSD from consOR1 bound to L-menthol MD simulations without miniGαs, and with miniGαs.

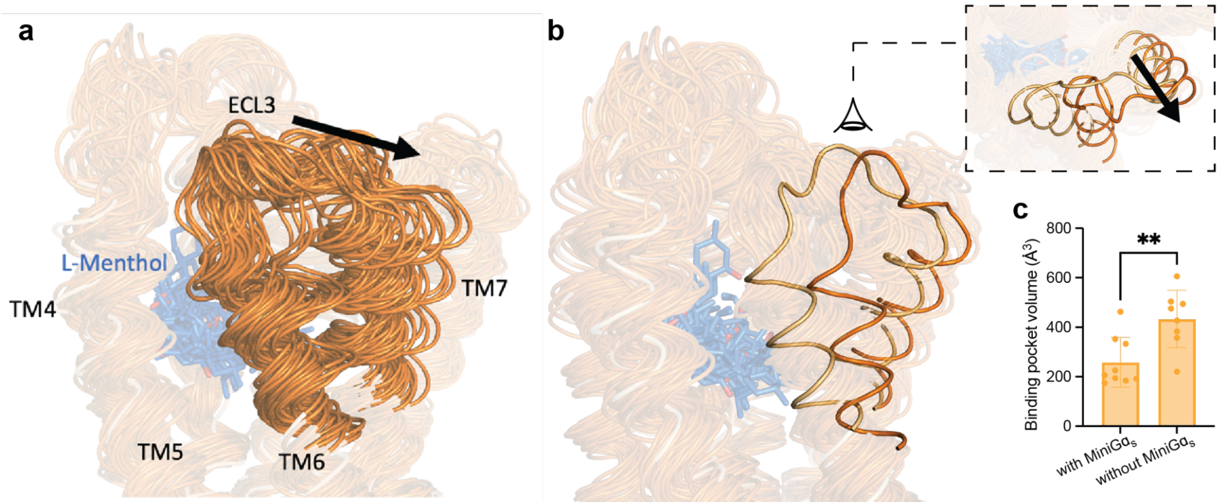


**Supplementary Figure 7. Extracellular motions during activation. a)** Sampling of conformations in consOR1 bound to menthol without mini Gαs MD simulations shows a motion of the extracellular parts of TM6 and TM7 as well as ECL3 away from the receptor bundle. **b)** Side view of extreme conformations, insert represent top view. **c)** consOR1 binding pocket volumes during MD simulations are bigger when the ligand-bound receptor is not bound to mini Gαs than when it is fully complexed.

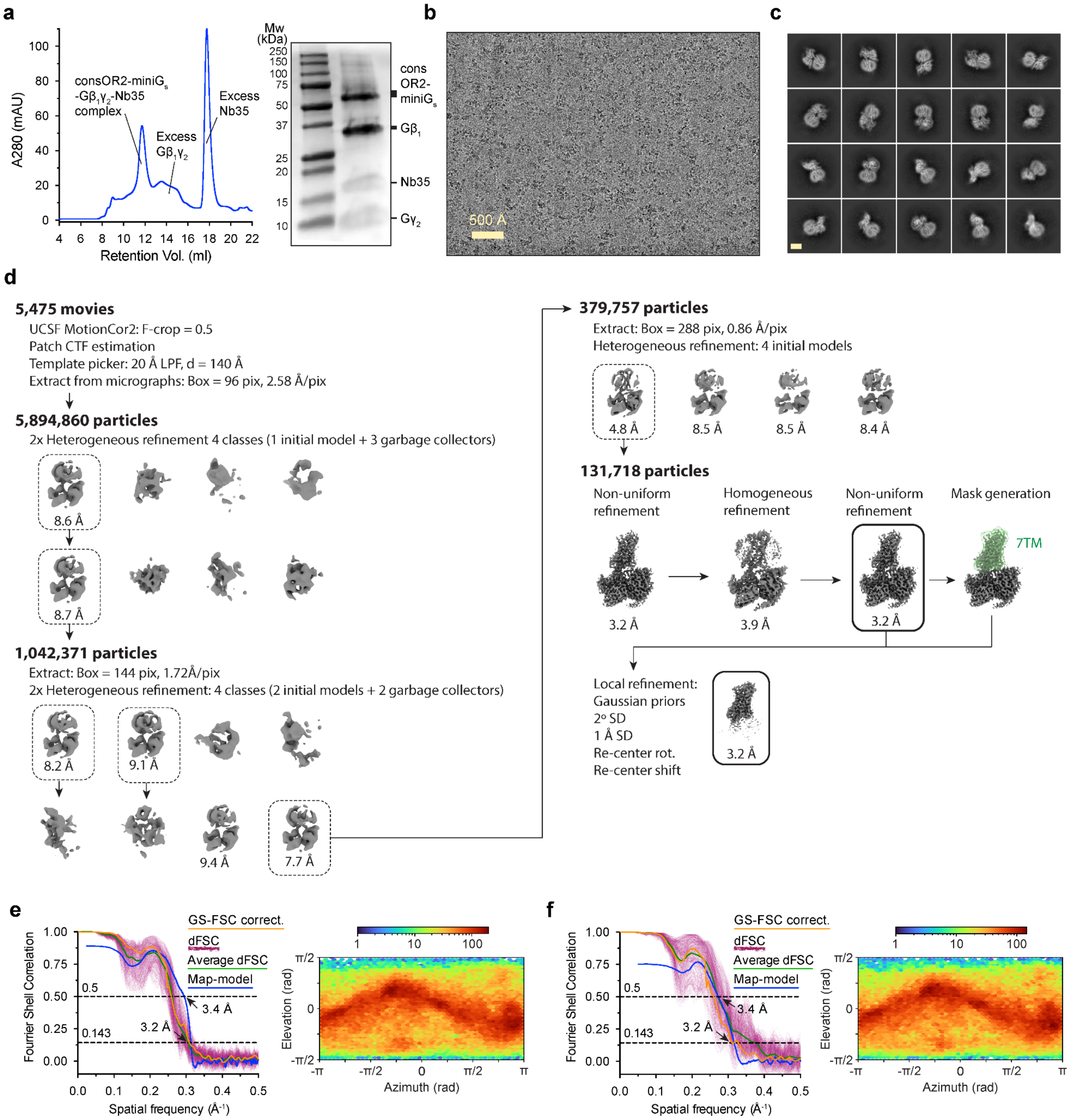


**Supplementary Figure 8. Cryo-EM data processing for consOR2-$G_s$. a)** Size exclusion chromatogram and SDS-PAGE of consOR2-$G_s$ sample. **b)** A representative cryo-EM micrograph from the curated consOR2-$G_s$ dataset (n = 5,475) obtained from a Titan Krios microscope. **c)** A subset of highly populated, reference-free 2D-class averages are shown. Scale bar is 50 Å. **d)** Schematic showing the image processing workflow for consOR2-$G_s$. Initial processing was performed using UCSF MotionCor2 and cryoSPARC. Particles were then transferred using the pyem script package[53] to RELION for alignment-free 3D classification. Finally, particles were processed in cryoSPARC using the non-uniform and local refinement tools. Dashed boxes indicate selected classes, and 3D volumes of classes and refinements are shown along with global Gold-standard Fourier Shell Coefficient (GSFSC) resolutions. **e, f)** Map validation for the consOR2-$G_s$ (**e**) globally refined, and (**f**) locally refined cryo-EM maps. Gold-standard Fourier shell correlation (FSC) curves are calculated in cryoSPARC, and shown together with directional FSC (dFSC) curves generated with dfsc.0.0.1.py as previously described[71]. Map-model correlations calculated in the Phenix suite are also shown. Arrows indicate map and map-model resolution estimates at 0.143 and 0.5 correlation respectively. Euler angle distributions calculated in cryoSPARC are also provided for each map.

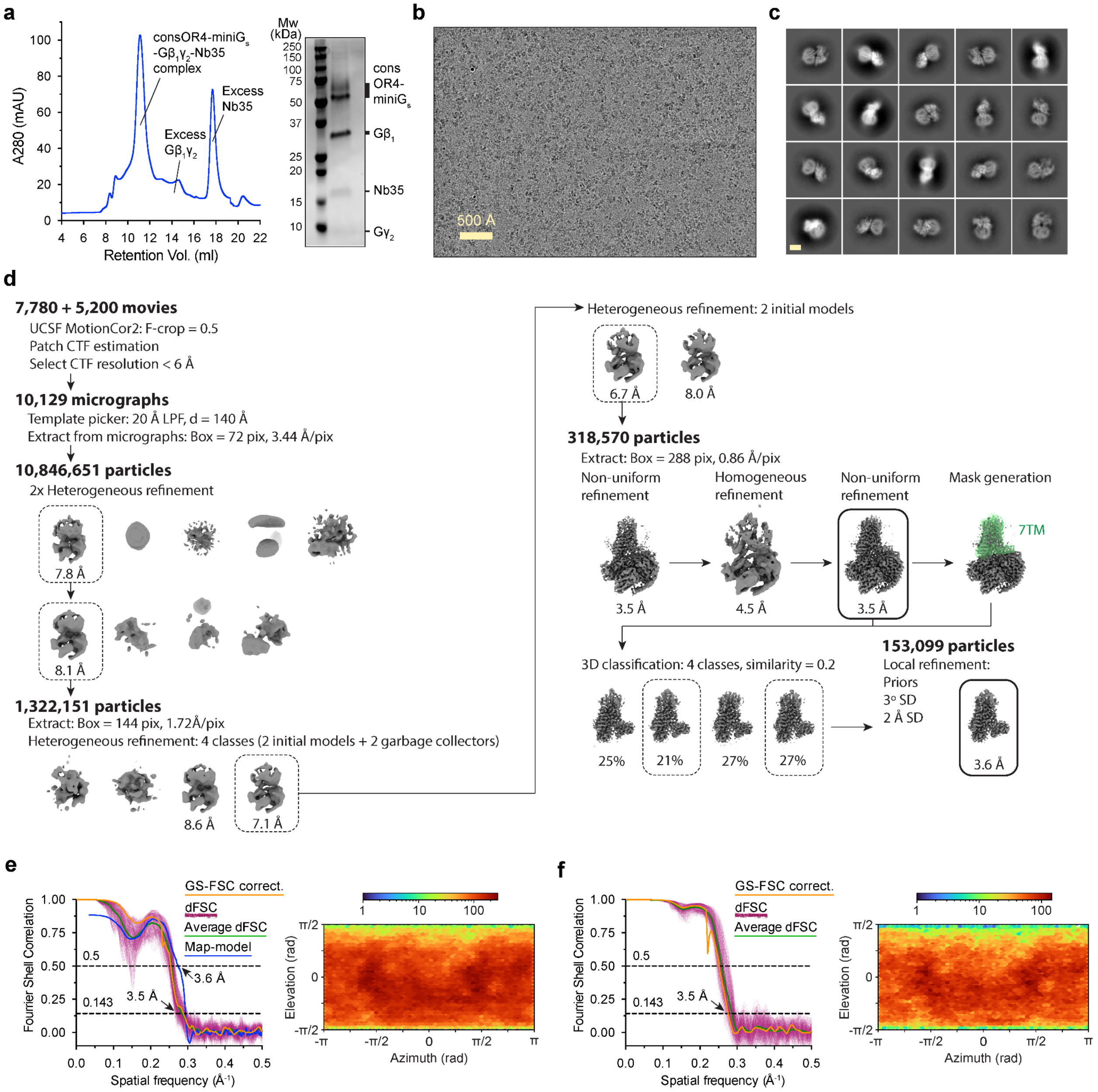


**Supplementary Figure 9. Cryo-EM data processing for consOR4-G$_s$. a)** Size exclusion chromatogram and SDS-PAGE of consOR4-G$_s$ sample. **b)** A representative cryo-EM micrograph from the curated consOR4-G$_s$ dataset (n = 10,129) obtained from a Titan Krios microscope. **c)** A subset of highly populated, reference-free 2D-class averages are shown. Scale bar is 50 Å. **d)** Schematic showing the image processing workflow for consOR4-G$_s$. Initial processing was performed using UCSF MotionCor2 and cryoSPARC. Particles were then transferred using the pyem script package[53] to RELION for alignment-free 3D classification. Finally, particles were processed in cryoSPARC using the non-uniform and local refinement tools. Dashed boxes indicate selected classes, and 3D volumes of classes and refinements are shown along with global Gold-standard Fourier Shell Coefficient (GSFSC) resolutions. **e, f)** Map validation for the consOR4-$G_s$ (**e**) globally refined, and (**f**) locally refined cryo-EM maps. Gold-standard Fourier shell correlation (FSC) curves are calculated in cryoSPARC, and shown together with directional FSC (dFSC) curves generated with dfsc.0.0.1.py as previously described[71]. Map-model correlations calculated in the Phenix suite are also shown. Arrows indicate map and map-model resolution estimates at 0.143 and 0.5 correlation respectively. Euler angle distributions calculated in cryoSPARC are also provided for each map.

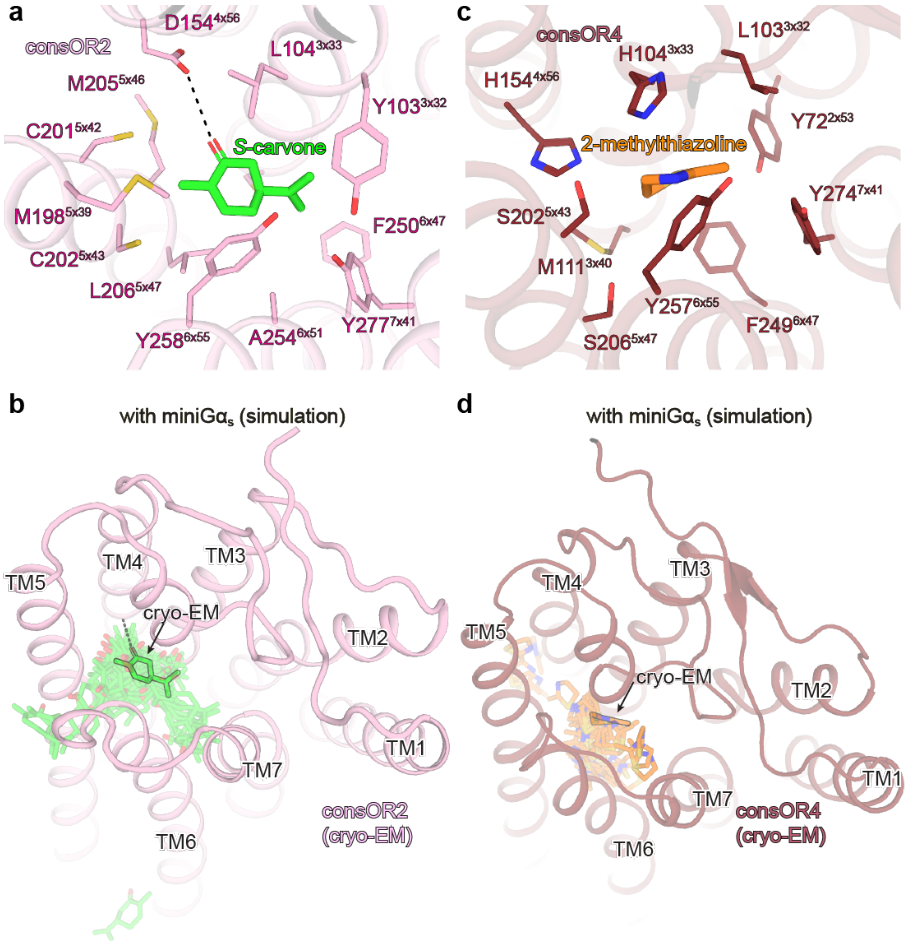


**Supplementary Figure 10. Odorant binding to consOR2 and consOR4. a)** The EM model of ligand in consOR2 with the ligand S-carvone contacting residues highlighted. **b)** The EM model of ligand in consOR4 with the ligand 2-methyl thiazoline contacting residues highlighted. In both panel a and b, the ligand contacting residues are found by utilizing UCSF Chimera Find Clashes/Contacts module with default parameter. **c)** The ligand positions during the consOR2 simulation. **d)** The ligand positions during the consOR4 simulation. In both panel c and d, the transparent ligands represent the simulation snapshots taken by skipping every 100 ns whereas the non-transparent ligand is the cryo-EM pose.

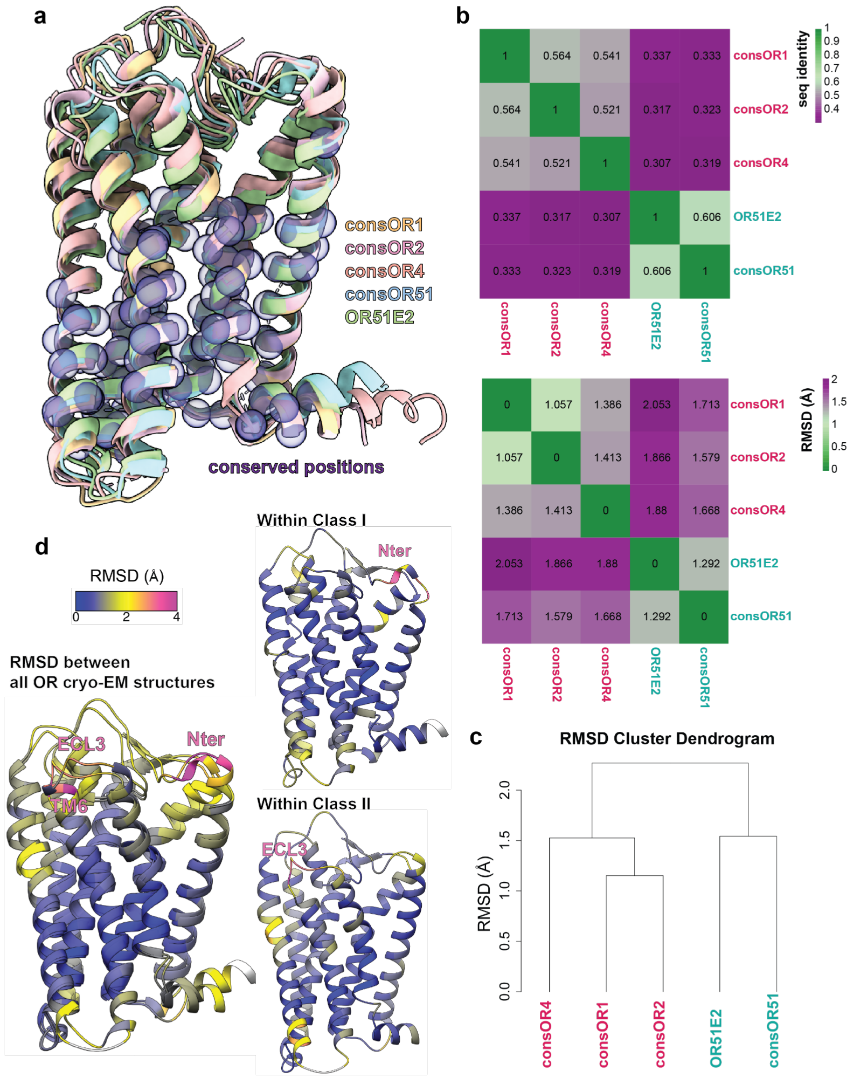


**Supplementary Figure 11. Comparison of all human OR cryo-EM structures. a)** Superimposition of all cryo-EM structures. The positions most conserved in terms of location are highlighted by purple dots. **b)** Heatmap of sequence identity and RMSD between the cryo-EM structures of consOR1, consOR2, consOR4 (red), consOR51 and OR51E2 (cyan). **c)** Cluster Dendrogram made by RMSD. Class II and Class I are clustered in two different groups. **d)** CryoEM structures colored by RMSD for all the structurally elucidated ORs (left, projected on consOR1 and consOR51), only Class I ORs (top right, consOR51 and OR51E2 projected on consOR51) and only Class II (bottom right, consOR1, consOR2, consOR4 projected on consOR1).

**Supplementary Table 1:** Sequence identity between consORs and native OR family members.

| Receptor | Maximum sequence identity (%) for native OR in family | Minimum sequence identity (%) for native OR in family | Average sequence identity (%) for all native ORs in family |
|---|---|---|---|
| consOR1 | 0.71 | 0.51 | 0.65 |
| consOR2 | 0.67 | 0.34 | 0.58 |
| consOR4 | 0.67 | 0.56 | 0.62 |
| consOR5 | 0.70 | 0.56 | 0.62 |
| consOR6 | 0.70 | 0.47 | 0.59 |
| consOR8 | 0.71 | 0.51 | 0.66 |
| consOR10 | 0.67 | 0.48 | 0.58 |
| consOR51 | 0.74 | 0.45 | 0.61 |
| consOR52 | 0.74 | 0.49 | 0.62 |

**Supplementary Table 2.** Cryo-EM data collection, refinement and validation statistics

| | consOR51-<br>$G_s$ complex | consOR1<br>L-menthol<br>$G_s$ complex | consOR2<br>S-carvone<br>$G_s$ complex | consOR4<br>2-MT<br>$G_s$ complex |
|---|---|---|---|---|
| EMDB: Full map | (EMDB-42786) | (EMDB-42789) | (EMDB-42791) | (EMDB-42817) |
| RCSB PDB: Model | (PDB 8UXV) | (PDB 8UXY) | (PDB 8UY0) | (PDB 8UYQ) |
| **Data collection and processing** | | | | |
| Magnification | 105,000 | 105,000 | 105,000 | 105,000 |
| Voltage (kV) | 300 | 300 | 300 | 300 |
| Electron exposure (e–/Å$^2$) | 50 | 50 | 50 | 50 |
| Defocus range (µm) | -2.0 to -0.8 | -2.1 to -1.0 | -2.1 to -1.0 | -2.1 to -1.0 |
| Pixel size (Å) | 0.873 (physical) | 0.86 (physical) | 0.86 (physical) | 0.856 (physical) |
| Symmetry imposed | *C1* | *C1* | *C1* | *C1* |
| Initial particle images (no.) | 6,275,495 | 15,310,482 | 5,894,860 | 10,846,651 |
| Final particle images (no.) | 204,513 | 212,883 | 131,718 | 318,570 |
| Map resolution (Å) (masked) | 3.2 | 3.3 | 3.2 | 3.5 |
| FSC threshold | 0.143 | 0.143 | 0.143 | 0.143 |
| **Refinement** | | | | |
| Initial model used (PDB code) | AlphaFold (consOR51)<br>7LJC (G protein)<br>3SN6 (Nb35) | AlphaFold (consOR1)<br>7LJC (G protein)<br>3SN6 (Nb35) | AlphaFold (consOR2)<br>7LJC (G protein)<br>3SN6 (Nb35) | AlphaFold (consOR4)<br>7LJC (G protein)<br>3SN6 (Nb35) |
| Model resolution (Å) | | | | |
| (unmasked/masked) | 3.4/3.2 | 3.9/4.0 | 3.6/3.4 | 3.8/3.6 |
| FSC threshold | 0.5 | 0.5 | 0.5 | 0.5 |
| Map sharpening *B* factor (Å$^2$) | -122 | -156 | -133 | -134 |
| Model composition | | | | |
| Non-hydrogen atoms | 8074 | 8153 | 8121 | 8240 |
| Protein residues | 1041 | 1035 | 1031 | 1042 |
| Ligands | 0 | 1 | 1 | 1 |
| *B* factors (Å$^2$) | | | | |
| Protein | 37.97 | 141.99 | 104.76 | 52.63 |
| Ligand | | 129.98 | 109.86 | 83.92 |
| R.m.s. deviations | | | | |
| Bond lengths (Å) | 0.003 | 0.003 | 0.004 | 0.003 |
| Bond angles (°) | 0.558 | 0.993 | 0.588 | 0.895 |
| Validation | | | | |
| MolProbity score | 1.22 | 1.04 | 1.23 | 1.35 |
| Clashscore | 3.11 | 2.09 | 4.50 | 3.82 |
| Poor rotamers (%) | 0 | 0.22 | 0 | 0 |
| CaBLAM outliers (%) | 0.99 | 1.29 | 0.40 | 1.19 |
| Ramachandran plot | | | | |
| Favored (%) | 97.37 | 97.75 | 98.12 | 96.97 |
| Allowed (%) | 2.63 | 2.25 | 1.88 | 3.03 |
| Disallowed (%) | 0 | 0 | 0 | 0 |

**Supplementary File 1**: Sequence identity between human ORs and the structurally elucidated consORs.
